# Replicating infant astrocyte behavior in the adult after brain injury improves outcomes

**DOI:** 10.1101/2020.05.14.096974

**Authors:** Leon Teo, Anthony G. Boghdadi, Jihane Homman-Ludiye, Iñaki Carril-Mundiñano, William C. Kwan, James A. Bourne

**Author notes:** Corresponding Author: James A. Bourne.

## Abstract

Infants and adults respond differently to brain injuries. Specifically, improved neuronal sparing along with reduced astrogliosis and glial scarring often observed earlier in life, likely contributes to improved long-term outcomes. Understanding the underlying mechanisms could enable the recapitulation of neuroprotective effects, observed in infants, to benefit adult patients after brain injuries. We reveal that in primates, Eph/ ephrin signaling contributes to age-dependent reactive astrocyte behavior. Ephrin-A5 expression on astrocytes was more protracted in adults, whereas ephrin-A1 was associated only with infant astrocytes. Furthermore, ephrin-A5 exacerbated major hallmarks of astrocyte reactivity via EphA2 and EphA4 receptors, which was subsequently alleviated by ephrin-A1. Rather than suppressing reactivity, ephrin-A1 signaling shifted astrocytes towards GAP43+ neuroprotection, accounting for improved neuronal sparing in infants. Reintroducing ephrin-A1 after middle-aged ischemic stroke significantly attenuated glial scarring, improved neuronal sparing and preserved circuitry. Therefore, beneficial infant mechanisms can be recapitulated in adults to improve outcomes after CNS injuries.

## Introduction

Beyond acute interventions, there are currently no neuroprotective drug strategies accessible to limit the extensive secondary damage that occurs in the sub-acute (days to weeks) stages after CNS injury, leading to poor outcomes. While improvements to the standard of care have increased survival rates, most patients continue to experience irrecoverable neurological or cognitive deficits due to damage in areas responsible for these modalities. A major impediment to neuronal sparing is the glial scar formed by reactive astrocytes at the injury site [1, 2]. Comprised of primarily reactive astrocytes, inflammatory cells and deposits of extracellular matrix chondroitin sulphate proteoglycan (CSPG), the glial scar has traditionally been associated with the inhibition of neuronal re-innervation into damaged regions [1].

While reactive astrocytes are key players that influence the severity of the glial scar and functional outcomes, under certain conditions, they exhibit neuroprotective functions that can be exploited to improve neuronal sparing and functional recovery after CNS injury [2, 3]. For example, limiting neuronal apoptosis, maintaining homeostasis and providing trophic support to surviving neurons [4–6]. Additionally, the capacity of reactive astrocytes to shift between states of reactivity, evidenced by changes in gene expression and function [7], demonstrates a high level of age- and environment-dependent plasticity [8–11] that drives the interplay between neuroprotective and neurotoxic roles. Indeed, evidence that the blockade of scar formation after spinal cord injury led to worse outcomes [12], indicates that the glial scar is, to an extent, required for CNS repair. Therefore, the identification and characterization of new functionally neuroprotective reactive astrocyte subpopulations may eventually birth novel strategies for therapeutic interventions after CNS injury and disease. Yet, the broad capacity of reactive astrocytes to be both neuroprotective and/or neurotoxic [13–15] confounds the therapeutic potential for clinical interventions.

However, an interesting phenomenon observed both clinically and in the laboratory setting is the age-dependent differences in the extent of secondary injury, whereby less severe functional deficits are observed after CNS injuries in infants, compared to adults [16–18]. This trend was highlighted in a previous study in which we demonstrated that the intrinsic and extrinsic regulators or astrocyte reactivity differed between infants and adults marmoset monkeys [10, 19]. This correlated to a smaller, more discrete chronic scarring which was more permissible to neuronal sparring after perinatal stroke compared to the equivalent injury in middle-aged adults [10]. Therefore, we hypothesized that manipulating reactive astrocyte activity at the injury site by reactivating infant-associated pathways has the potential to attenuate scarring and improve outcomes following an adult brain injury.

Our study focused on the family of glycosylphosphatidylinositol (GPI)-linked ephrin-A ligands and the EphA mammalian receptor tyrosine kinases. A wealth of evidence has linked these guidance cues with the pathophysiology and pathogenesis of CNS injury. Specifically, Eph/ ephrin signaling is a crucial regulator of astrocyte reactivity in rodents [20–24], nonhuman primates [10, 25] and human [26]. Furthermore, the Eph/ ephrins have been the target of a number of preclinical treatment studies [27, 28]. The significant transcriptional differences between rodent and human astrocytes [29], as well as the more protracted pathophysiological time course in primates [10, 19] were accounted for by using a clinically relevant model of perinatal and middle-aged focal ischemia in the marmoset monkey’s primary visual cortex (V1) of [19].

## Results

### Age-dependent upregulation of ephrins in response to ischemic brain injury

We first examined the expression of members of the EphA/ ephrin-A family in the infant and adult nonhuman primate (marmoset monkey) primary visual cortex (V1) after a vasoconstrictor-induced focal ischemic injury (Fig. 1A-D). Most striking was a significant elevation in the expression of ephrin-A1 in the infant but not adult post-ischemic brain at 7 days post-injury (DPI; Fig. 1E, F, J; Supplementary Fig. 1). Both age groups revealed increased expression of ephrin-A5 (Fig. 1E, G, K; Supplementary Fig. 1), previously associated with increased astrocyte reactivity via EphA4 receptor activation [23, 25] but which was sustained only in the adult at 7 DPI. The upregulation of ephrin-A1 and -A5 were associated with GFAP+ reactive astrocytes, proximal to the injury site at their respective peak of upregulation (Fig. 1H-M; Supplementary Fig. 2, 3). For EphA receptors, EphA1 was not detectable in all conditions tested. We confirmed the increase of EphA4, a modulator of astrocyte reactivity, in the chronic period after focal ischemia, remaining elevated in both conditions at 21 DPI (Fig. 1E). EphA2 expression was transiently diminished 1-week post injury in infants but not adult (Fig. 1E). Finally, a sustained decrease in EphA7 expression was detected in infants only from 7 to 21 DPI (Fig. 1E). EphA7 expression is primarily associated with neurons in both rodents and primates [30]. The reduction of EphA7 in infants without a similar occurrence in adults, where neuronal loss is more severe [10], most likely indicates neuronal downregulation of EphA7 expression in infants.

**Figure 1.**
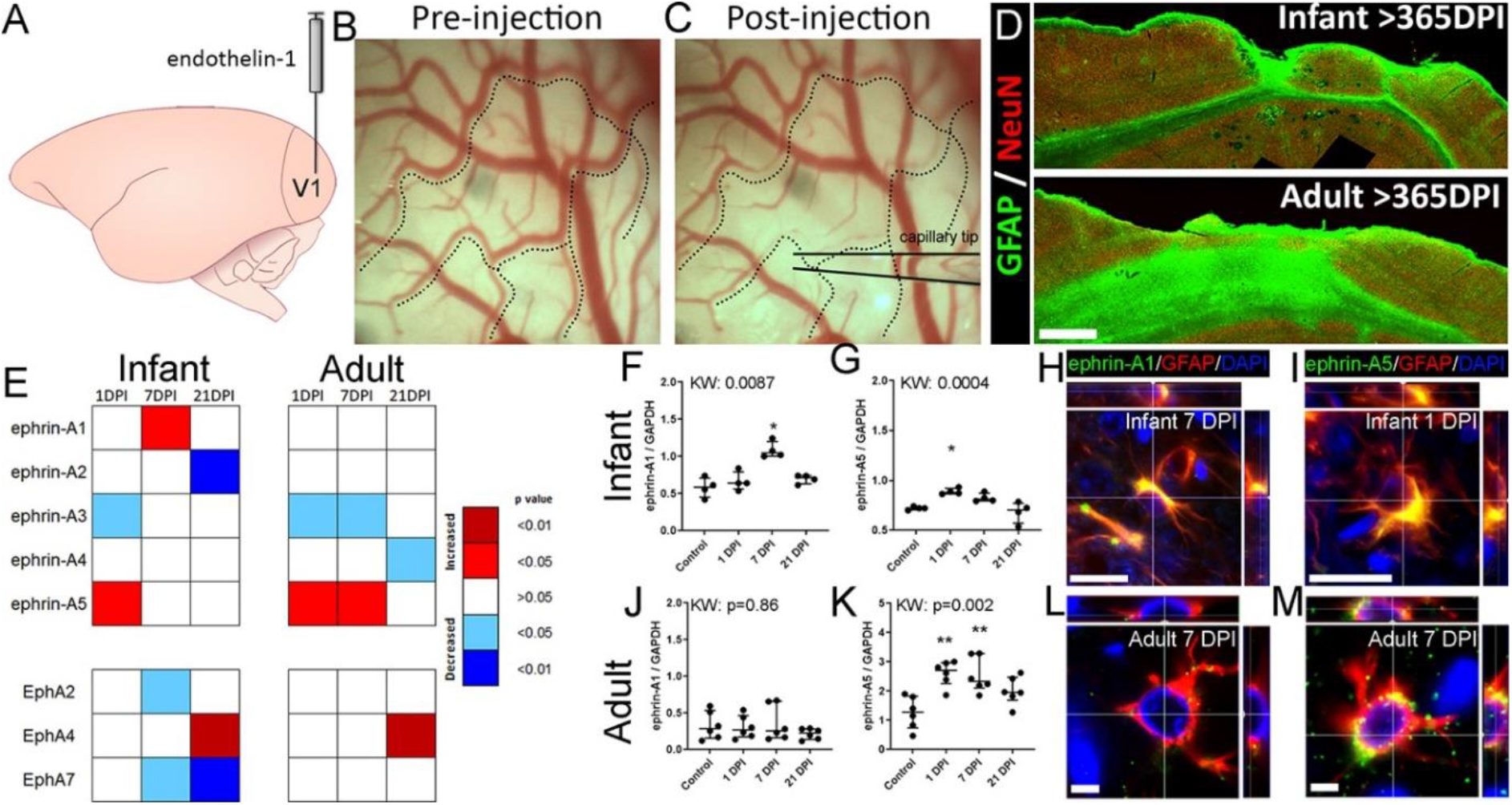
Ephrin-As’ expression on reactive astrocytes differ following infant versus adult focal ischemia to the marmoset V1. (A) Cartoon of the marmoset neocortex with the primary visual cortex (V1) indicated. The vasoconstrictor endothelin-1 was injected proximal to the calcarine branch of the posterior cerebral artery. (B, C) Photomicrographs of V1 surface taken during the focal ischemia induction surgery. The vasoconstrictor endothelin-1 was administered intracortically proximal to the distal branches of the calcarine artery (dotted lines) (B), resulting in vaso-occlusion (C). (D) GFAP and NeuN immunolabeled sections >1 year after lesion induction surgery in the infant and adult marmoset V1. GFAP+ glial scarring is less severe after infant injury compared to adult ^[4]^. (E) Heatmap summarizing the Eph/ephrin expression via quantitative immunoblots screen at 1, 7 and 21 days after infant or adult focal ischemia. Ephrin-A1 expression was increased in infants (F) but not adults (J). Ephrin-A5 expression was increased post-injury in infants and adults with more prolonged upregulation in adults (G, K). The upregulation of ephrin-A1 and -A5 were associated with GFAP+ reactive astrocytes in the injured infant (G, I) and adult (L, M) V1 during the peak of their respective upregulation. Scale: (D) 0.5mm, (H-M) 22μm. (A) recreated under the Creative Commons Attribution-Share Alike 4.0 International license.

**Figure 2.**
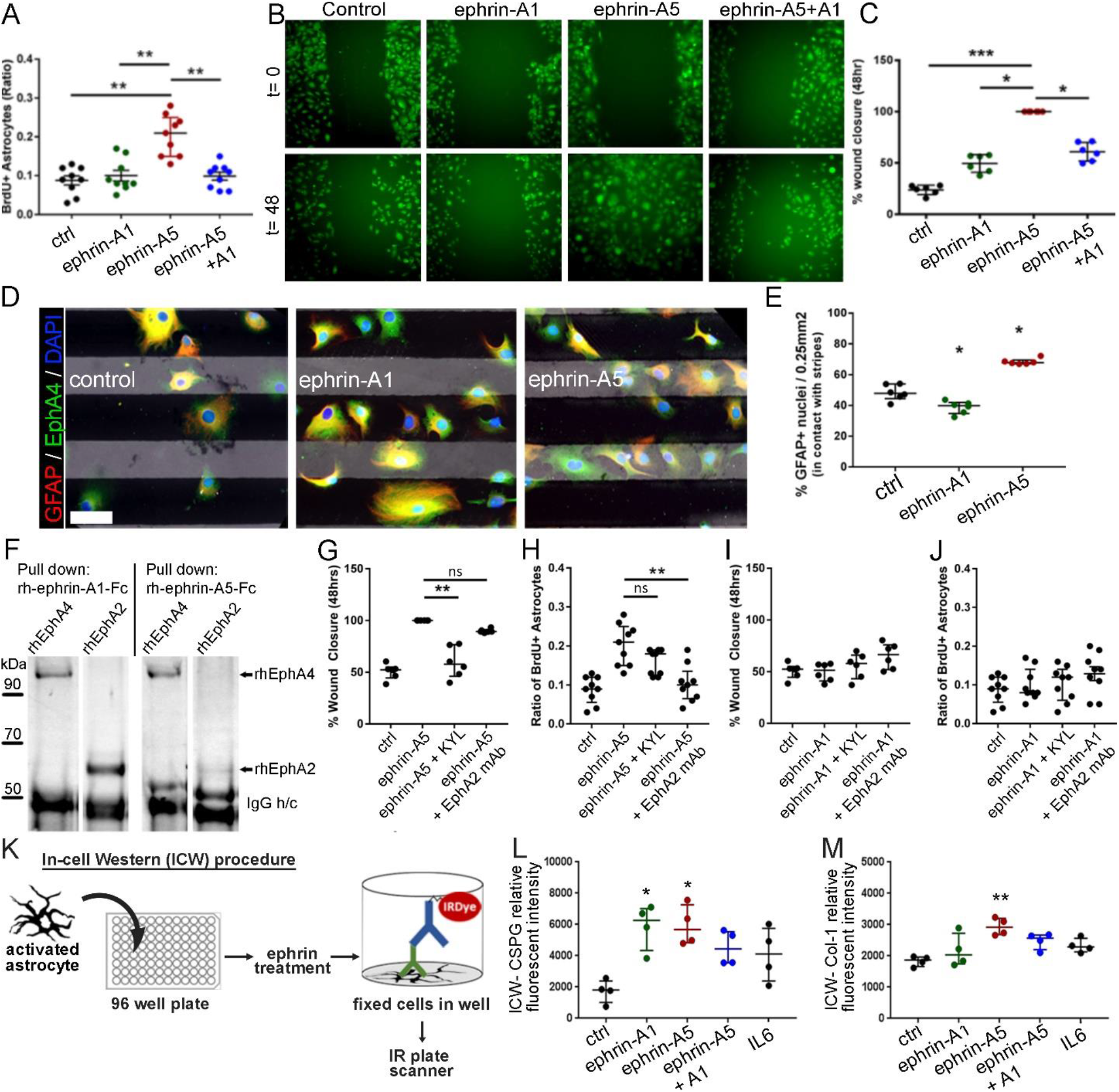
EphrinA1 signaling attenuates ephrin-A5 induced astrocyte reactivity. (A) Ephrin-A5 signalling significantly increased astrocyte proliferation, while ephrin-A1 signalling did not. Addition of ephrin-A1 significantly attenuated ephrin-A5 induced astrocyte proliferation to baseline levels. (B) Photomicrographs of astrocyte wound closure assays at t=0h and t=48h. (C) Ephrin-A5 signalling significantly increased astrocyte wound closure at t=48h while ephrin-A1 signalling did not. Addition of ephrin-A1 significantly attenuated ephrin-A5 induced astrocyte wound closure. (D, E) In stripe assays, ephrin-A5 stripes induced attraction while ephrin-A1 stripes induced repulsion on astrocytes migration. (F) Pull down assays confirmed that both ephrin-A1 and ephrin-A5 binds EphA2 and EphA4 receptors. (G) Ephrin-A5 induced astrocyte wound closure was significantly attenuated to baseline levels through KYL induced EphA4 receptor blockade while EphA2 blockade did not. (H) EphA2 blockade significantly reduced ephrin-A5 induced astrocyte proliferation but EphA4 blockade did not. (I, J) Neither KYL induced EphA4 blockade, nor mAB induced EphA2 blockade significantly had any significant effect on ephrin-A1 treated wound closure or proliferation assays. (K) Illustration demonstrating in-cell-western protocol using IL6-activated astrocytes. (L) Both ephrin-A1 and ephrin-A5 signalling significantly increased astrocyte CSPG secretion, which was significantly reduced by combined treatment of ephrin-A1 and –A5. (M) Ephrin-A5 signalling significantly increased astrocyte collagen-1 expression but ephrin-A1 did not. Addition of ephrin-A1 significantly attenuated ephrin-A5 induced collagen-1 expression to baseline levels. * p<0.05; ** p<0.01; *** p<0.001. Scale bar: (D) 50μm.

**Figure 3.**
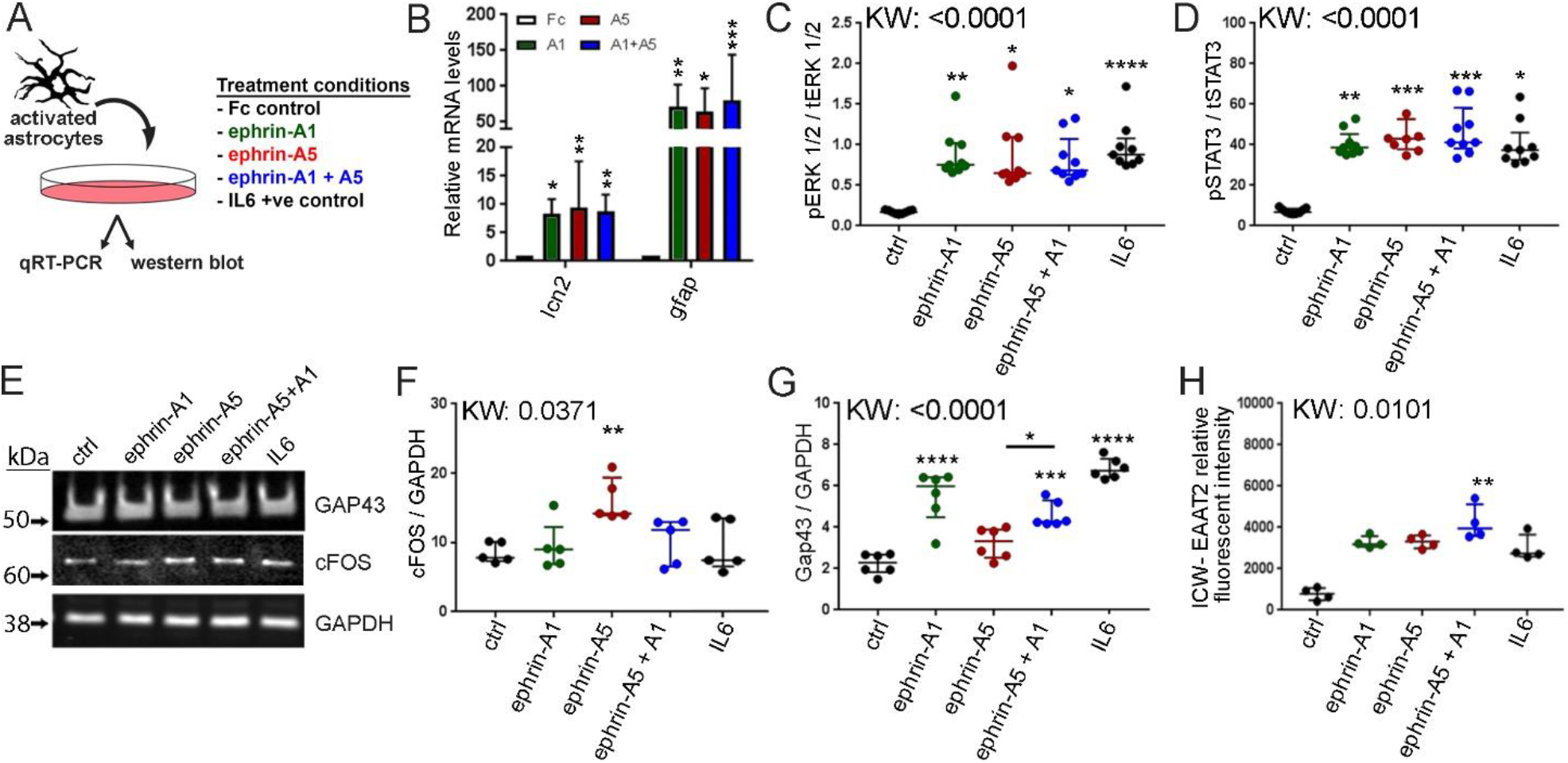
Ephrin-A1 signaling increases expression of reactive astrocyte markers associated with neuroprotection. (A) Cartoon illustrating the experimental procedure. (b-d) Both ephrin-A1 and –A5 signaling increased the expression of pan-reactive astrocyte markers on activates astrocytes: lcn2 and GFAP (B), as well as increased ERK1/2 (C) and STAT3 (D) phosphorylation, all of which were not affected by combined ephrin-A1 + -A5 treatment. (E, F) Ephrin-A5 signaling induced upregulation of cFos on reactive astrocytes, associated with increased proliferation and neuroinflammation, but ephrin-A1 did not. Addition of ephrin-A1 significantly attenuated ephrin-A5 induced reactive astrocyte cFOS expression back down to baseline levels. (E, G) Ephrin-A1 signaling induced upregulation of GAP43 on reactive astrocytes, associated with neuroprotection against glutamate excitotoxicity, but ephrin-A5 did not. The addition of ephrin-A1 to ephrin-A5 conditions successfully induced the expression of GAP43 expression to ephrin-A1 levels, corresponding to an upregulation of the excitatory amino acid transporter 2 (EAAT2; H). * p<0.05; ** p<0.01; *** p<0.001; ****p<0.0001.

### Ephrin-A1 and -A5 signaling induces divergent astrocyte function

Next, we determined if ephrin signaling contributes to the age-dependent astrocyte responses [10] by investigating the functional effects of ephrin-A1 and -A5 on several hallmarks of astrocyte reactivity in vitro. To emphasize clinical translatability, we used primary derived human cortical astrocytes in our in vivo assays. Reactive astrocyte proliferation is a key characteristic of reactivity after injury [10, 31]. We examined the effects of ephrin-A1 and –A5 signaling on astrocyte proliferation via thymidine analogue incorporation. We confirmed that ephrin-A5 treatment significantly increased astrocyte proliferation (Fig. 2A) [25, 32], while ephrin-A1 did not. This correlates with our in vivo data demonstrating that the timing of ephrin-A5 upregulation (Fig. 1E) directly corresponds to the respective peaks of reactive astrocyte proliferation post-ischemia in infants and adults [10]. Centripetal migration of reactive astrocytes to the injury site is another hallmark of astrocyte reactivity [33–35]. In scratch wound experiments [36], We established that ephrin-A5 significantly increased astrocyte wound closure (Fig. 2B, C, Supplementary video 1) [25], while ephrin-A1 did not. As Eph/ ephrin signaling typically guides cell-migration [37], we tested the effects of ephrin-A1 and –A5 signaling on astrocyte guidance in a stripe assay. Ephrin-A5 stripes induced cell attraction, while ephrin-A1 stripes induced cell repulsion (Fig. 2D, E), contradicting prior evidence of ephrin-A5 induced cell-repulsion [38]. This is likely attributed to negligible Src (c-Src; rodent homolog: pp60c-Src) expression in human astrocytes [29, 30], compared to mice (Supplementary Fig. 4), which reverses ephrin-A5 induced cell guidance [39]. To assess the translatability and representativeness between our marmoset and human *in vitro* astrocyte systems, we repeated the scratch wound and stripe assays using marmoset astrocytes [40; Supplementary Fig. 5]. Our results demonstrate that ephrin-A1 and -A5 treatment induces identical effects on marmoset astrocyte centripetal migration (Supplementary Fig. 5B, C, Supplementary video 2) and cell guidance (Supplementary Fig. 5D, E) as observed in human astrocytes. These findings make evident the capacity for ephrin-A5 signaling to induce several hallmarks of astrocyte reactivity, specifically proliferation and centripetal migration of reactive astrocytes to the injury site, while ephrin-A1 signaling does not.

To determine the specific EphA receptor/s involved, we selected EphA2 and EphA4 as receptor candidates due to their high affinity with ephrin-A1 and –A5 [41] (Fig. 2F). Expression of EphA2 and EphA4 was first confirmed on GFAP+ reactive astrocytes in vivo, proximal to the injury site (Supplementary Fig. 6, 7). Next, we established that EphA4 blockade (KYL) decreases ephrin-A5 induced wound closure, while EphA2 blockade (EphA2 mAb) did not (Fig. 2G). In proliferation assays, EphA2 blockade significantly reduced ephrin-A5 induced astrocyte proliferation but EphA4 blockade did not (Fig. 2H). Blockade of EphA2 and EphA4 had no effect on ephrin-A1 treated assays (Fig. 2I, J). Thus, EphA2 and EphA4 activation by ephrin-A5 increases astrocyte motility and cell proliferation, respectively. These results provide greater insights into how the various hallmarks of astrocyte reactivity can be influenced by ephrin-As/ EphAs signaling.

**Figure. 4.**
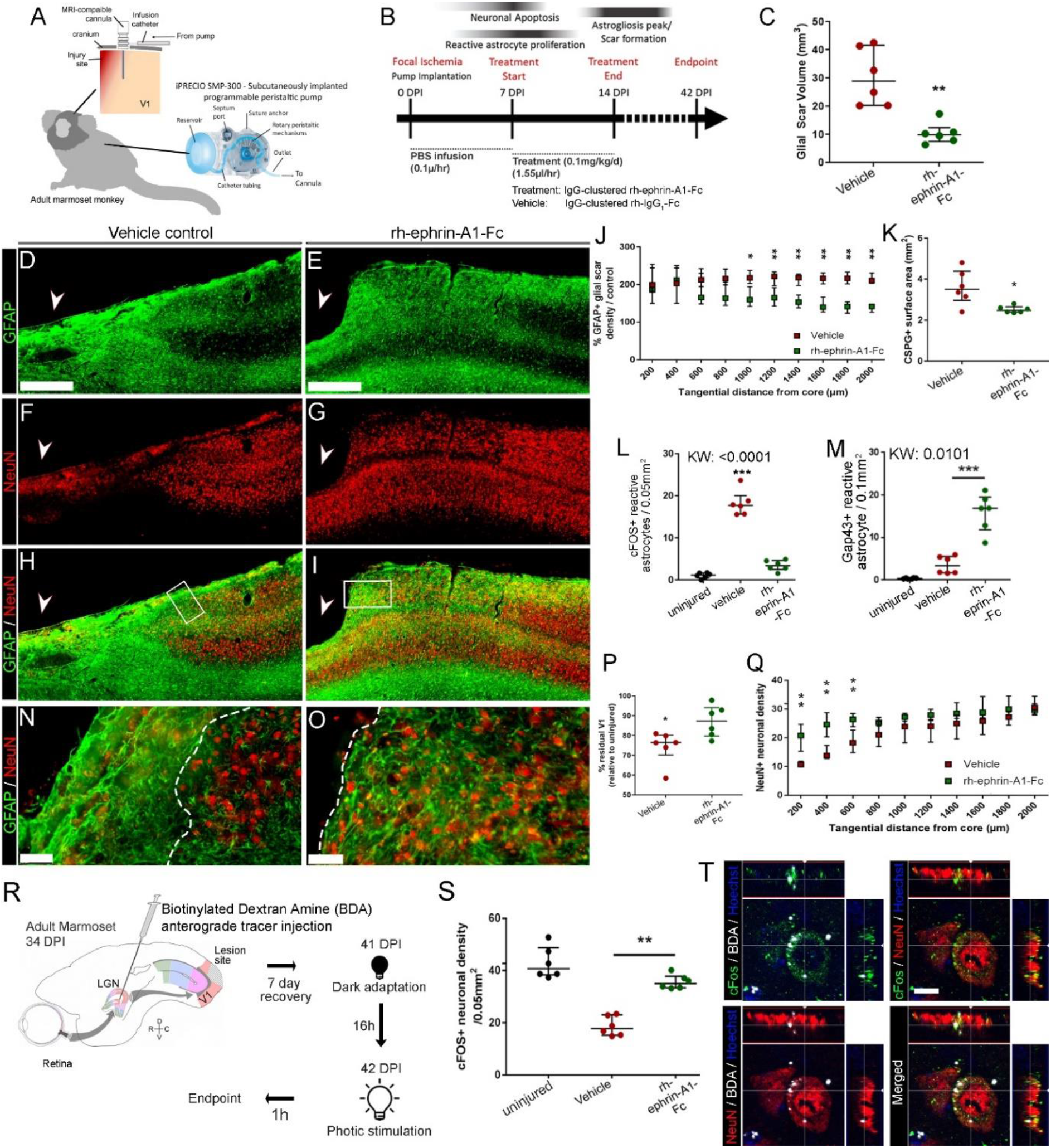
Rh-ephrin-A1-Fc infusion attenuates glial scarring and increases sparing of visual responsive neurones after ischemic injury in the adult marmoset V1. (A) Cartoon illustrating the *in vivo* rh-ephrinA1-Fc treatment experiment. (B) *In vivo* treatment timeline. A subcutaneously implanted peristaltic pump commenced the infusion of rh-ephrin-A1-Fc or vehicle (IgG_1_-Fc in sterile PBS) at 7 days post injury (DPI), corresponding to the peak in astrocyte proliferation post injury. (C) Rh-ephrin-A1-Fc infusion significantly reduced GFAP+ glial scar volume, compared to vehicle controls. (D-I) GFAP and NeuN immunofluorescent labelled parasagittal sections of vehicle and rh-ephrin-A1-Fc treated V1 comprising the lesion core (arrow) and peri-infarct area. Bounding boxes in (H, I) denotes enlarged region in (N, O). (J) GFAP+ reactive astrocyte optical density was significantly reduced in rh-ephrin-A1-Fc treated cohorts proximal to the infarct core. (k) Reduction in glial scarring and GFAP+ reactive astrocyte density was correlated to a reduction in CSPG deposition proximal to the lesion core. (L) Rh-ephrin-A1-Fc treatment significantly reduced the subpopulation of proliferative / neuroinflammatory cFOS+/GFAP+ reactive astrocyte proximal to the lesion core. (M) Rh-ephrin-A1-Fc infusion induced a significant increase in a subpopulation of neuroprotective GAP43+/GFAP+ reactive astrocyte within 1mm of the lesion core. (P) Rh-ephrin-A1-Fc infusion significantly increased NeuN+ residual V1 volume, compared to vehicle controls. (Q) Increased NeuN+ optical density was detected proximal to the lesion core after rh-ephrin-A1-Fc treatment, indicative of neuronal sparing. (N) The glial scar can be clearly delineated as a GFAP+ rich region devoid of neurones (dotted lines). (O) Rh-ephrin-A1-Fc infusion resulted in an increase in NeuN+ neuronal sparing within the GFAP+ reactive astrocyte-rich glial scar region, extending into the edge of the lesion core. (R) Illustration demonstrating the procedure for neural tracer injection followed by dark-adaptation/ photic stimulation protocol to visualise cFOS+ light responsive neurons in V1. (S) rh-ephrin-A1-Fc infusion significantly increased the sparing of cFOS+/NeuN+ light responsive neurons, which retained inputs from the lateral geniculate nucleus (T) proximal to (within 1mm) the lesion core. * p<0.05; ** p<0.01. Scale bar: (D-I) 0.5mm, (N-O) 50μm, (t) 10μm.

### Ephrin-A1 attenuates ephrin-A5-induced astrocyte reactivity *in vitro*

To determine whether ephrin-A1 treatment could potentially attenuate astrogliosis, we performed combined treatments of ephrin-A1 and –A5 on astrocytes *in vitro* to mimic the upregulation of ephrin-A5 at 7 DPI (Fig. 1E). Ephrin-A1 treatment significantly attenuated the effects of ephrin-A5 induced proliferation (Fig. 2A) and wound closure (Fig. 2B, C, Supplementary video 1) back to baseline levels, most likely through disruption of ephrin-A5 signaling. We next evaluated the effects of ephrin-A1 and –A5 signaling on reactive astrocyte scar formation using in-cell-western assays (Fig. 2K) to measure CSPG and collagen-1 (col-1) levels, which correlates with more severe glial scar outcomes [8, 42]. Both ephrin-A1 and –A5 alone significantly elevated CSPG secretion (Fig. 2L), consistent with scar formation at both ages [10]. However, only ephrin-A5 treatment significantly increased collagen-1 expression (Fig. 2M), required for secondary transformation into scar-forming astrocytes [8]. Importantly, ephrin-A1 treatment successfully attenuated ephrin-A5-induced elevation of both CSPG (Fig. 2L) and collagen-1 (Fig. 2M) back to baseline levels. These results provide an explanation as to how ephrin-A1 signaling tempers ephrin-A5-induced hallmarks of astrocyte reactivity, contributing to a less severe chronic glial scar outcome (Fig. 1D) [10], after CNS injury in infants.

### Ephrin-A1 nor -A5 signaling suppresses astrocyte reactivity

To further investigate how rh-ephrin-A1-fc treatment influences astrocyte reactivity and/or function, cultured astrocytes were IL-6 activated [8, 43, 44], and specific markers of astrocyte reactivity were measured following ephrin-A treatment (Fig. 3A). While quantitative real-time PCR analysis revealed increased levels of pan-reactive markers lipocalin2 and GFAP following treatment with the ephrin-As, compared to negative controls (Fc; Fig. 3B), no significant differences were detected between groups. Similarly, Erk1/2 (Fig. 3C) and STAT3 (Fig.3D) phosphorylation, crucial modulators of astrocyte activation [43, 45] and scar formation [46], were increased compared to negative controls with no significant difference between treatment groups. Increased STAT3 activation, an important regulator of scar formation [46], in all treatment conditions indicates that the scar-forming capacity is not reduced by ephrin-A treatment. Altogether, this suggests that the attenuation of ephrin-A5-induced reactive astrocyte responses by ephrin-A1 were not consequences of suppressed astrocyte reactivity.

### Ephrin-A1-induced phenotypic changes to reactive astrocytes

The function of a particular subpopulation of reactive astrocyte (neurotoxic vs. neuroprotective) is directly influenced by a host of intrinsic and extrinsic cues present after injury [8, 10, 13, 47, 48]. This plasticity underpins the complex functional and phenotypic heterogeneity of reactive astrocytes after injury. As ephrin-A1 treatment does not suppress astrocyte reactivity, we postulate that ephrin-A1 signaling triggers functional changes on reactive astrocytes that confers beneficial outcomes. We examined if ephrin-A1 or -A5 signaling on astrocytes are associated with the “A1” neurotoxic or “A2” neuroprotective subpopulations [13] previously described in rodents [13, 48]. We determined that neither ephrin-A1 nor –A5 signaling, individually or combined, was responsible for- or sufficient to induce conversion of reactive astrocytes into either “A1” or “A2” phenotypes (Supplementary Fig. 8A, B). Additionally, in neither of these two subpopulations was ephrin-A1 or –A5 upregulated [13] above baseline levels (Supplementary Fig. 8C). This is either due to interspecies differences between rodent and primates, or a consequence of the diverse states of reactivity that drives reactive astrocytes’ response to injury [7]. However, we did observe that ephrin-A5 treatment significantly increased *cFOS* expression on reactive astrocytes (Fig. 3E, F) *in vitro*, associated with increased proliferation [49] and pathological neuroinflammation [50]. In contrast, ephrin-A1 treatment significantly increased GAP43 expression on reactive astrocytes (Fig. 3E, G), while ephrin-A5 did not. GAP43+ reactive astrocytes are a distinct neuroprotective subpopulation capable of protecting against glutamate excitotoxicity [9]. To better represent the *in vivo* environment in infants, we combined treatment with ephrin-A1 and –A5, which resulted in the suppression of ephrin-A5-induced cFOS upregulation back to baseline levels (Fig. 3E, F). Ephrin-A1 treatment also significantly increased the expression of GAP43 expression compared to the baseline levels in ephrin-A5 treated cohort (Fig. 3E, G). The increased expression of GAP43 with combined ephrin-A1 and -A5 also corresponded with the upregulation of excitatory amino acid transporter 2 (EAAT2; Fig. 3H), required for the GAP43-dependent neuroprotection against glutamate excitotoxicity [9]. From these *in vitro* studies, it appears that the discrete scar outcomes in infants are not due to a reduction in astrocyte reactivity but most likely a consequence of ephrin-A1-dependent phenotypic changes, which promote more neuroprotective states of reactivity.

### Ephrin-A1 treatment attenuates chronic scarring *in vivo* after injury

With significant *in vitro* evidence indicating that ephrin-A1 signaling can override ephrin-A5-induced reactive astrocyte functions and induce neuroprotection in response to injury, we further investigated its therapeutic potential *in* vivo (Fig. 4A). To establish whether ephrin-A1 contributes to the reduced glial scar observed after an infant brain injury [10], we administered a local intracerebral infusion of rh-ephrin-A1-Fc into the post-ischemic adult marmoset V1. Treatment was continuously infused from 7-14DPI, corresponding to the period of peak reactive astrocyte proliferation post-injury [10] (Fig. 4B). Glial scar severity was assessed at the chronic 42 DPI endpoint. Our results demonstrate distinct glial scarring in both treatment cohorts, consistent with increased STAT3 activation *in vitro* (Fig. 3D). However, volumetric analysis revealed a marked decrease in total GFAP+ glial scar volume in rh-ephrin-A1-Fc treated V1 (Fig. 4C, E, I), compared to vehicle controls (0.1mg/kg/day IgG-clustered recombinant human IgG1-Fc; Fig. 4C, D, H) was detected. This reduction in glial scarring was attributed to a significant decrease in GFAP+ reactive astrocyte density observed 1,000-2,000μm from the infarct core (Fig. 4J), corresponding directly to reduced chondroitin sulfate proteoglycan (CSPG) deposition (Fig. 4K, Supplementary Fig. 9). The attenuation of glial scarring observed in adults following rh-ephrin-A1-fc treatment mimicked the discrete and less severe glia scar previously observed in infants following a brain injury (Fig. 1D) [10, 19]. Additionally, consistent with our *in vivo* studies, we detected a large subpopulation GFAP+/ cFOS+ (proliferative [49]/ neuroinflammatory [50]) reactive astrocytes proximal to the injury site in vehicle controls (Supplementary Fig. 10, 11), but were significantly reduced following rh-ephrin-A1-fc infusion (Fig. 4l; Supplementary Fig. 10, 11). A significant increase in GFAP+/ GAP43+ (neuroprotective [9]) reactive astrocytes was also detected within 1mm proximal to the injury site following rh-ephrin-A1-fc infusion in the post-ischemic adult V1 (Fig. 4m; Supplementary Fig. 12) compared to vehicle controls. We further infer that rh-ephrin-A1-fc signaling had no effect on microglia activity *in vivo* based on subthreshold expression levels of both receptors on microglia [29] (Supplementary Fig. 8D).

### Ephrin-A1 treatment preserves neuronal circuitry and function *in vivo* after injury

We further examined the extent of neuronal sparing after infusions of rh-ephrin-A1-fc (Fig. 4A, B) in the post-ischemic adult primary visual cortex (V1). Volumetric analyses revealed an increase in the percentage of NeuN+ residual V1 volume in rh-ephrin-A1-fc treated cohorts (Fig. 4G, I, P) compared to vehicle controls (Fig. 4F, H, P), indicative of neuronal sparing. This was supported by increased NeuN+ neuronal density proximal to the infarct core (Fig. 4H, I, Q), corresponding to dense GFAP+ scar regions (Fig. 4D-I). In vehicle controls (Fig. 4N), the dense glial scar was devoid of neurons, yet following rh-ephrin-A1-fc treatment (Fig. 4O) there was a significantly greater number of NeuN+ neurons within the GFAP+ glial scar, and juxtaposed the terminal edge of the infarct core.

These findings indicate that while ephrin-A1 treatment does not completely ameliorate the formation of a glial scar in the injured adult brain, it promotes an environment that is more permissible to neuronal survival, as previously observed in infants [10]. To determine if spared neurons proximal to the infarct core retained their afferent connectivity and were still part of a functional circuit, we performed MRI-guided injection of an anterograde tracer into the lateral geniculate nucleus (LGN; Fig. 4R) followed by a dark adaptation/ photic stimulation protocol [51] to induce expression of the immediate-early gene, *cFOS*, a marker of active/ functioning visual neurons. A greater density of light-responsive neurons was detected within 1mm proximal to the infarct core in rh-ephrin-A1-fc treated V1 (Fig. 4S, Supplementary Fig. 10), which were recipient of inputs from the LGN (Fig. 4T) compared to vehicle controls. These results confirm that the ephrin-A1 treated reactive astrocytes that form the glial scar are capable of sparing neurons and their function following injury.

## Discussion

The greater capacity of the infant brain to respond to injury is dependent on multiple factors. These include reduced myelination and ECM deposition, as well as the presence of growth factors during postnatal development that likely contributes to greater neuronal survivability. Here we reveal that age-dependent ephrin-A1 treatment on reactive astrocytes post-injury leads to reduced glial scar severity observed in infants following a brain injury [10], contributing to improved neuronal sparing. This is supported by prior evidence that reactive astrocyte-associated ephrin-A1 signaling contributes to functional recovery after spinal cord injury in rodents [21]. Additionally, ephrin-A1 signaling can be recapitulated following brain injury in adults to attenuate scarring and improve outcomes. Due to the interspecies differences highlighted here and in previous work [10, 19, 52, 53], this necessitates the need to examine mechanisms of injury in model systems which more closely resembles that of the human, especially if clinical translation is the predominant purpose. For example, ephrin-A1 and -A5 treatment elicited identical outcomes in both human and marmoset astrocytes *in vitro*, which differed from that observed in rodents [38]. Our conclusions highlight that ephrin-A1 signaling on reactive astrocytes [21] is required for improved functional recovery after CNS injury. Importantly the rh-ephrin-A1-fc infusion tempered rather than abrogated glial scarring in adults, providing further evidence that glial scarring is a necessary component and prerequisite to repair [12]. Together, this study provides compelling evidence that targeted recapitulation of the enhanced repair mechanisms and plasticity [42] inherent during early life is an effective strategy for the future therapeutic developments aimed at improving outcomes after CNS injuries in adults.

Astrogliosis and glial scarring have traditionally been recognized as major barriers to repair and regeneration in the mammalian CNS [1], inhibiting the re-innervation of neurons to re-establish damaged circuitry. A long-held belief was that astrogliosis is synonymous with glial scarring and is, therefore, categorically detrimental to repair. However, recent studies have demonstrated that while reactive astrocytes can exacerbate secondary neuronal degeneration [54], they also provide crucial neuroprotection post-injury (e.g. limit neuronal apoptosis, maintain homeostasis, debris clearance and provide trophic support to surviving neurons [15]). Furthermore, the blockade of scar formation after spinal cord injury led to worse outcomes [12], indicating that the glial scar is, to an extent, required for CNS repair. This is directly reflected in our findings, wherein topical administration of rh-ephrin-A1-Fc *in vivo* after focal ischemia attenuated but did not abrogate scar formation. Rather, rh-ephrin-A1-Fc treatment induced significant phenotypic changes that tempered the traditionally compact, dense glial scar, resulting in improved neuronal sparing. Moreover, the capacity of reactive astrocytes to shift between states of reactivity — each of which with discrete gene expression identity and function [7] — demonstrates a high level of age- and environment-dependent plasticity [8–11] that drives the interplay between neuroprotective and neurotoxic roles. Indeed, our results indicate that rh-ephrin-A1-Fc treatment may promote reactive astrocyte-mediated neuroprotection [44] against glutamate excitotoxicity through a GAP43/NFκB-dependent increase in astrocyte-specific EAAT2 [9]. Rh-ephrin-A1-Fc treatment also reduced cFos-associated reactive astrocyte proliferation [55, 56], which may have additional implications on neuroinflammation and neurotoxicity [56].

A point of interest is the diverging guidance induced by ephrin-A5/ EphA4 signaling on primate vs. rodent cells [38]. While there is a case to be made for interspecies differences, indicated by differences in Src expression, it should also be noted that ephrin-A5/ EphA4 signaling can induce both attraction and repulsion effects on axonal pathfinding [57]. The promiscuity and functional diversity of Eph/ ephrin interactions raises the possibility for the involvement of other Eph receptors on reactive astrocytes in our treatment paradigm. Due to the lack of a comprehensive Eph/ ephrin expression profile for reactive astrocytes after injury, we cannot rule out the involvement of other Eph receptors, in addition to the possibility of ephrin-A-mediated reverse signaling on reactive astrocytes. However, our data suggests that ephrin-A1 signaling disrupts ephrin-A5/ EphA2 (ephrin-A1’s cognate receptor) [58] and EphA4 (high-affinity receptor) [41] -dependent astrocyte reactivity. This is consistent with the known roles of EphA2 and EphA4 as modulators of cell migration [59–61], cell-cycle [25, 62, 63] and astrocyte reactivity [10, 23, 25]. Specifically, the involvement of ephrin-A5/ EphA4 signaling on reactive astrocytes, which promotes proliferation and motility, via the mitogen-activated protein kinase (MAPK) and Rho-dependent pathways [25, 64, 65]. Interestingly, the co-expression of ligand and receptors on reactive astrocytes indicates a level of cell-contact mediated communication that may contribute to the local regulation of astrogliosis. This Eph/ephrin mediated local regulation could contribute to increased local proliferation and cell-cell attraction, accounting for the dense accumulation of reactive astrocytes observed in adults. Additionally, while less is known about its effect on reactive astrocytes, EphA2 activation on endothelial cells and various tumor cells negatively regulates cell proliferation [66–68], likely through c-Myc suppression and stabilization of the cyclin-dependent kinase inhibitor p27/KIP1 [67]. Competitive inhibition of EphA4 signaling by rh-ephrin-A1-Fc (vs. endogenous ephrin-A5) on reactive astrocytes could account for the attenuation of scarring observed. This is supported by prior evidence that ephrin-A1 and -A5 bind with similar high affinity to EphA4 (ephrin-A1: Kd = 0.27; ephrin-A5: Kd = 0.25) [41]. Furthermore, EphA2 activation is increased following EphA4 blockade [32], accounting for the reduction in reactive astrocyte proliferation and consequent scar density after treatment. Future development of ephrin-A1 as a therapeutic target will need to fully elucidate the downstream effects of ephrin-A1 on reactive astrocytes, its direct and indirect effects on neuroprotection, as well as the exact Eph receptors involved. Furthermore, we acknowledge that the IgG-mediated clustering of rh-ephrin-A1-Fc (required to enable signal propagation [69]) results in a large protein complex that is not directly translatable for clinical applications and is therefore, a limitation of our approach. This can be remedied through drug design to enable the presentation of nanoscale, multivalent ephrin-A1 mimetics/ analogues, e.g.: via biopolymer conjugation [70], to facilitate clinical translation.

Although improvements to the standard of care and diagnostics tools have increased survival rates, treatment options remain limited to the brief therapeutic windows following the immediate insult. For example in ischemic stroke: clot-busting drugs (<4.5hrs) [71], mechanical thrombectomy (<6-24hrs) [72] or decompressive craniotomies (<48hrs) [73]. Therefore, any therapy that can reduce neuronal loss and enhance recovery in the extended period (days-to-weeks) beyond the current limits has the potential to markedly improve patient outcomes and reduce the burden of disease. The administration of ephrin-A1 treatment at 7 days post-injury, corresponding to the peak of secondary neuronal injury in primates [19] has the potential to fulfill this unmet need in CNS therapeutics. This is most significant in cases where early intervention treatment is not possible, thereby ensuring that patients receive greater access to CNS therapeutics. Importantly, since comparable mechanisms are at play on reactive astrocytes following most types of CNS injuries, the application of this strategy can broadly encompass disorders where astrogliosis and glial scarring are major barriers to neuronal sparing and functional recovery, including traumatic brain [74, 75] or spinal cord injuries [12, 76, 77] and strokes [10, 78, 79].

Neurons have historically been the center of attention in neuroscience research [80] that sought to: 1. Investigate the normal function of the CNS; 2. Elucidate the consequences of injury and disease: and, 3. Develop treatment strategies for alleviating pathologies. In this context, the potential for neuroprotection by reactive astrocytes has been relatively neglected. Whilst therapeutic interventions aimed at rescuing neuronal function after CNS injury is crucial, equally important is harnessing the supportive and neuroprotective function of astrocytes to establish a stable and permissive milieu to maximize neuronal survival and functional sparing. What has become clear recently is that astrocytes play roles far beyond that of maintaining homeostasis and inducing scar formation after injury. One major leap forward has been the breakthrough surrounding the functional heterogeneity of astrocytes, which includes multi-species gene expression profiling [52] and the identification and characterization of discrete subpopulations of reactive astrocytes [8, 9, 13, 81–83]. These studies have only begun to untangle the profound age [10, 11] and environment [8] dependent plasticity that drives neurotoxic vs. neuroprotective functions after CNS injury. This has therefore led to a gradual shift in research perspective aimed at better understanding the causal, regulatory and adaptive roles that astrocytes play in the CNS [80]. This is most evident in fields of CNS trauma, stroke, epilepsy and neurodegenerative diseases where the pathogenesis and pathophysiology of disease are in part driven by astrocyte reactivity.

## Materials and Methods

### Animals

36 outbred marmoset monkeys (*Callithrix jacchus*) aged postnatal (P) 14 (n=15), and >5 years (n=21; middle-aged) were used in this study. Gender was not a criterion in the selection of the animals and no siblings were used. Animals were housed in family groups (12:12 hrs light/dark cycle, temperature 31°C, humidity 65%). Experiments were conducted according to the Australian Code of Practice for the Care and Use of Animals for Scientific Purposes and were approved by the Monash University Animal Ethics Committee. Animals were obtained from the National Nonhuman Primate Breeding and Research Facility (Monash University).

### Cell culture

Primary human cortical astrocytes purified from 17-18 weeks gestation human fetuses of indeterminate sexes were used in this experiment (Lonza). Astrocytes were expanded using the astrocyte growth medium bulletkit according to the manufacturer’s instructions at 37°C, 5% CO_2_. Astrocytes were expanded in serum-containing medium. While serum is excluded from the brain under normal physiological conditions, the focal ischemia induced in our model does result in the extravasation of blood, including serum proteins into the brain, detectable between 1-21 days post ischemia [19] (within the treatment period). This is concomitant with observations in rodent and human [84] after stroke. While we acknowledge that the use of serum-containing media may not be appropriate in studies relating to normal / quiescent astrocyte function [85], studies relating to astrocyte reactivity under pathological conditions require separate considerations. Serum-treated astrocytes were used in studies primarily focused on CNS pathologies and reactive astrocyte function [8, 43, 44, 46]. All experiments were performed between passages 2-3, due to the downregulation of GFAP in subsequent passages. Astrocytes were activated according to protocol adapted from [8, 43, 44]. Briefly, were plated in DMEM 10% FBS, 1X pen/strep and 1X GlutaMAX and recovered for 3 days, equilibrated under serum-free conditions for 24 hours before stimulation with 50ng/ml IL6 (Abcam) and 200ng/ml soluble IL6 receptor (Abcam) for 16-18 hours prior to ephrin treatment. For receptor blockade experiments, cells were pre-blocked using EphA4 antagonist peptide (KYL; Tocris) or EphA2 monoclonal antibody (Abcam; 10μg/ml) 1h prior to as well as during ephrin treatment to maintain continuous blockade.

Marmoset astrocytes were derived from neurospheres generated from excised marmoset V1 tissue at PD14 using protocols published previously [40]. Astrocytes were generated by passaging neurospheres 3 times to maximize astrocyte yield, followed by 1 week in NeuroCult NSA with differentiation supplement (StemCell Technologies) and further expanded in Neurocult NSA with proliferation supplement (StemCell Technologies; 10ng/ml rhFGF-2, 20ng/ml rhEGF) and incubated at 37°C, 5% CO_2_. Wound-closure and stripe assays using marmoset astrocytes were performed using similar protocols as described for human astrocytes.

### Induction of focal ischemic injury

Induction of focal ischemic injury to the infant and adult marmoset V1 was performed by vasoconstrictor-mediated vascular occlusion of the calcarine branch of the posterior cerebral artery (PCAca), as detailed previously [10, 19]. Briefly, following anesthesia (Alfaxalone 5mg/kg; adults only; maintained with inspired isoflurane 0.5-4%), a craniotomy and dural thinning was performed, followed by intracortical injections of endothelin-1 (ET-1; 0.1 μl / 30s pulse at 30s intervals, totaling ~0.5 μL for per site for infants-4 sites and ~0.7 μL for adults-7 sites) proximal to the calcarine artery, which supplies operculum V1. Upon completion, the craniotomy was replaced, secured with tissue adhesive (Vetbond; 3M) and the skin sutured closed. Naive animals were used as uninjured controls.

### Peristaltic pump and intracranial cannula implantation

For adult animals undergoing rh-ephrin-A1-fc or vehicle treatments, a cutaneous incision was performed under the right scapula, parallel to the midline prior to the closure of the craniotomy. A subcutaneous pocket was created under the left scapula using blunted hemostats and a sterile pre-programmed micro-infusion pump (iPrecio; SMP-300) was inserted into the subcutaneous pocket and secured using anchoring sutures. The attached tubing was threaded subcutaneously towards the cranial incision. An intra-cerebral cannula (3280PM; 30G needle; Plastics One) was installed on the excised bone flap using bone cement and adjusted to 1.5mm needle penetration depth. The cannula was implanted directly outside of a 1mm radius exclusion zone from lesion epicenter, located using the stereotaxic micromanipulator to enable the infusion of treatment solution in the surviving parenchyma, as determined in our previous study [19]. The tubing was attached to the cannula port and the bone flap was reaffixed to the cranium with orthopedic cement.

### Infusion protocol

Each peristaltic pump was activated, pre-loaded with sterile phosphate buffered saline (PBS; 0.01M) and pre-programmed before implantation. Pumps were programmed to initiate administration of PBS (0.1μL/hr) to the injury site throughout a 7-day recovery period to prevent blockage of the cannula. Treatment was designed to begin at 7 DPI, corresponding to the peak of reactive astrocyte proliferation as well as the peak of secondary neuronal apoptosis from secondary injury [10, 19]. At this time, the remaining PBS in the reservoir withdrawn through the port septum. The pump reservoir was refilled with treatment solution comprising pre-clustered rh-ephrin-A1-fc (see ephrin clustering) or vehicle control (IgG clustered human IgG_1_ fc) in sterile PBS. A 10μL/hr (2.5h) flush was included in the infusion protocol to clear any remaining PBS in the tubing and cannula. Treatment infusion proceeded over 7 days (7-14 DPI) at a rate of 1.55μL/hr to administer a final dose of ~0.1mg/kg/day. A constant infusion of a relatively high dose was performed to increase availability of the treatment peptide and mitigate risk of potential degradation. The pump reservoir was refilled with treatment solution after 3.5 days using the port septum. Pumps were removed at the conclusion of treatment protocol and animals were recovered for 42 days to allow for chronic glial scar formation.

### MRI-guided tracer injection and photic stimulation

Injection of the biotinylated dextran amine (BDA 10, 000) anterograde tracer was performed at 34DPI, as previously described [86]. Briefly, animals were anesthetized and positioned in a custom-designed MRI-compatible stereotaxic frame. A midline incision was performed followed by a craniotomy. Fiducial markers (2mM gadolinium solution; Gadoteridol) were attached at the midline using tissue adhesive before MRI scanning. Fast-spin echo T2-weighted sequence was performed to demarcate brain areas and determine stereotaxic coordinates. Parameters: repetition time (TR)/echo time (TE) = 6,000/40 ms, echo train length = 4, field of view (FOV) = 38.4 × 38.4 mm^2^, acquisition matrix = 192 × 192, 100 coronal slices, slice thickness = 0.4 mm, and signal averages = 4. Using the MRI visualization software OsiriX, the fiducial markers in the frame and on the cranium were aligned to confer an accurate representation of elevation/ azimuth and roll to calculate the stereotaxic coordinates of the LGN. Injections of 400nl 10% BDA 10,000 were performed using a 33-gauge Neurosyringe (Hamilton) with a beveled tip at a rate of 180nl/min. Following completion, the craniotomy was reaffixed, skin resutured and animals were recovered for 7 days to allow for tracer transport. At 41DPI, animals were dark-adapted for at least 16h followed by 1h photic stimulation immediately prior to euthanasia.

### Fresh tissue collection

At the end of the designated recovery period, marmosets designated for ephrins screen (n=3/ cohort) were administered an overdose of pentobarbitone sodium (100 mg.kg^−1^; intraperitoneal). Following apnea, cerebral tissues were recovered and dissected under aseptic conditions in sterile ice-cold phosphate buffered saline (PBS; 0.1M; pH 7.2). Hemispheres were separated and the occipital lobe was dissected at the level of the diencephalon and bisected coronally. Caudal portions, encompassing V1 and the infarct core were collected and snap-frozen in liquid nitrogen.

### Perfusion and fixed tissue processing

Animals were euthanized as above. Following apnea, infants were transcardially perfused with warm 0.1M heparinized phosphate buffered saline (PBS, pH 7.2) containing 0.1% sodium nitrite, and adults with heparinized PBS, followed by 4% paraformaldehyde (PFA). Brains were dissected, post-fixed and cryoprotected, as outlined in our previous study [10, 87]. Following separation of the hemispheres, each hemisphere was bisected coronally at the start of the caudal pole of the diencephalon and frozen in liquid nitrogen at −40°C. The occipital block which encompasses V1 was cryosectioned in the parasagittal plane at −20°C.

### Western Blot

Proteins for western blotting were purified using TRIzol-LS reagent (Life). Snap-frozen tissue was homogenized in 750μL per 50-100μg-tissue weight and processed according to the established protocols [88]. Cells were lysed using RIPA buffer (Thermo) and processed according to the manufacturer’s instructions. Protein concentrations were determined through a modified Bradford protein assay (Bio-Rad). Following protein separation on 4-12% bis-tris polyacrylamide gel electrophoresis, proteins were blotted onto polyvinylidene difluoride membranes (PVDF; Millipore) and blocked in blocking buffer (1:1; Li-Cor) in 0.1M PBS for 30 mins. Incubation with primary antibodies (Supplementary Table 1) in blocking buffer diluted 1:1 with 0.1% PBS-Tween was performed overnight at 4°C. Following washes, membranes were incubated with secondary antibodies (Supplementary Table 2) in 0.1% PBS-Tween containing 0.02% SDS for 1 hr at room temperature. Protein bands were visualized using the Odyssey CLx infrared imaging system (Li-Cor).

### Eph/ephrin binding assay

Recombinant human (rh) ephrin-A1-fc or rh-ephrin-A5-fc (5μg) were conjugated to protein A Dynabeads (Life) for 10 min at room temperature. Following 0.1% PBS-TW washes, ephrin-conjugated beads were treated with purified rh-EphA4 or rh-EphA2 extracellular domains (5μg; R&D Systems) overnight at 4°C with agitation to determine binding. Following 0.1% PBS-TW washes, conjugates were eluted with glycine (pH 2.0), electrophoretically separated and blotted as described above.

## *In vitro* experiments

### Ephrin ligand clustering

Clustering of Eph receptors and ligands is required for signal propagation [69]. Rh-ephrin-A1-fc (R&D systems; 6417-A1-050) and -A5-fc (R&D systems; 374-EA-200) ligands (67μg/μl) were pre-clustered through incubation with anti-human-fc antibody (54 μg/μl; Sigma; I12136) at 37°C for 2 hrs and adjusted to the final concentration (10 μg/μl) using the appropriate culture medium. Negative controls were performed for all experiments using antibody-clustered rh-IgG_1_-fc.

### Wound-closure assay

8-well chamber slides (Lab-Tek) were coated overnight with 1mg/ml poly-L-lysine solution at 37°C, rinsed and coated with 1mg/ml laminin in MEM for 2 hrs at 37°C. Astrocytes were plated at high density (5×10^4^ cells/well), as described above, and allowed to recover overnight. Cells were incubated for 45 mins with 10μM CellTracker Green CMFD (Life) diluted in the fresh culture medium. A longitudinal scratch was then performed across the astrocyte monolayer using a sterile 200μL pipette tip. Wells were rinsed once and incubated with ephrin treatment solution and transferred to a humidity-controlled chamber (37°C, 5% CO_2_) fitted on a Leica AXF600 LX inverted fluorescent microscope and multi-positional time-lapsed imaging was performed over 48 hrs at 30 min intervals for temporal qualitative and quantitative analyses.

### Proliferation assay

Chambers slides were coated overnight with 1mg/mL poly-L-lysine and laminin as described. Astrocytes were plated at low density (2.625 ×10^4^ cells/well) and allowed to recover for 3 days. Cells were subsequently equilibrated and synchronized under serum-free conditions for 24h prior to ephrin treatment in the presence of 0.2μM BrdU and subsequently fixed in 4% PFA.

### Stripe assay

13mm diameter glass coverslips were coated with poly-L-lysine, dried and positioned coated side down on a silicone matrix with 50μm wide alternating grooves 50μm apart (obtained from Dr Martin Bastmeyer, Karlsruhe University, Germany) [89]. Pre-clustered ephrin ligands or vehicle were injected into the matrix and incubated for 2 hrs at room temperature. Following a PBS flush, a solution of 2% BSA-Alexa488 (Invitrogen) in PBS was injected into the matrix and incubated for 2h to visualize the stripes. Following a second PBS flush, coverslips were placed in 24-well plates and coated with laminin as described. Astrocytes were plated (4 ×10^4^ cells) on each coverslip and incubated for 18 hrs followed by fixation in 4% PFA.

### In cell western (ICW)

Astrocytes were plated on coated 96 well plates at 2 ×10^4^ cells/ well and recovered for 1-week to encourage extracellular matrix production before IL6 stimulation. Following 24h ephrin stimulation, treatment solution was replaced with DMEM with 1X pen/strep and 1X GlutaMAX for a further 2 days before fixation in 4% PFA. Fixed cells were washed in PBS, blocked with 10% serum in 0.1% PBS-TW and incubated in primary antibodies (Supplementary Table 1) at 4°C overnight. Following washes, cells were treated with secondary antibodies (Supplementary Table 2) in 0.1% PBS-Tween for 1h at room temperature. Following washes, cells were treated with CellTag700 for normalization. Wells were air-dried before signal visualization using the Odyssey CLx infrared imaging system (Li-Cor). Standard densitometry was performed and data were expressed as protein fluorescent intensity ratio over total cell numbers.

### qRT-PCR

Astrocytes were plated on 6 well plates at high density 5 ×10^5^/ well and stimulated as described. Following 24h ephrin treatment, cells were rinsed and lysed using TRIzol-LS reagent (Life) and RNA purified according to established protocol [88]. Total RNA was quantified using a Nanodrop spectrophotometer (OD_260_) and cDNA generated from 2μg total RNA using random hexamers primers (SuperScript IIIl; Life). qRT-PCR was performed using LightCycler 480 SYBR Green Master (Roche) according to the manufacturer’s specifications. Results were quantified and expressed as 2^^−(ΔΔct)^ normalized to HPRT1 expression. The primers used in this study are listed in (Supplementary Table 3).

### Immunohistochemistry, immunocytochemistry and histology

For immunohistochemistry, free-floating sections were washed in PBS before treatment with 0.3% hydrogen peroxide and 50% methanol in PBS for 20 min to inactivate endogenous peroxidases. Sections were pre-blocked in a solution of 10% normal goat serum in PBS + 0.3% Triton X-100 (TX; Sigma) before incubation with primary antibodies overnight at 4°C (Supplementary Table 1). Sections were rinsed in 0.1% PBS-Tween and incubated with biotinylated secondary antibodies (Supplementary Table 2) for 1 hr at room temperature. Sections were then treated with streptavidin-horseradish peroxidase conjugate (GE Healthcare; 1:200) and visualized via a metal-enhanced chromogen, 3,3’-diaminobenzidine (DAB; Sigma).

Immunofluorescent antigen detection was conducted with similar procedures to those above, except steps including and after secondary antibody incubation was performed in the dark. Following overnight incubation at 4°C with primary antibodies in blocking solution (Supplementary Table 1). Sections were then rinsed in 0.1% PBS-Tween before incubation with secondary antibodies (Supplementary Table 2). Detection of biotinylated dextran amine (BDA) tracer was performed by incubating the sections with streptavidin Alexa Fluor at 1:1000 at room temperature for 1h. Sections were also treated with either Hoechst 333258 or 4’,6-diamidino-2-phenylindole (DAPI) nuclei stains. Fluorescent Immunocytochemistry (ICC) experiments were performed similarly. Incubation of coverslips with secondary antibodies were performed for 30 mins, at room temperature.

### Histology

Nissl-substance (cresyl violet) staining was performed to aid in the anatomical and laminar demarcation of V1 in the infant and adult marmoset brain based on methods previously described [19].

### Microscopy

Standard brightfield and epifluorescent microscopic examination of processed sections was conducted using Zeiss SteREO discovery (V20) and Axio Imager Z1 microscopes (Zeiss). Images were obtained with a Zeiss Axiocam (HR rev3) using Axiovision software (V4.8.1.0; Zeiss). Tile-scanning for were performed using the Hamamatsu Nanozoomer, Zeiss Axioscan.Z1 and the Olympus VS120 slide scanners. All photomicrographs were cropped and resized using Photoshop (CC 2015.0.1, Adobe). Necessary image manipulations (eg. Resolution adjustment, pseudocoloring, brightness/contrast adjustments, image cropping, photomerging and the addition of legends, arrows and bounding boxes) were performed to aid in thr analysis and improve the quality of final images.

### Qualitative and Quantitative analyses

Multiplex western blot (WB) densitometry and ICW fluorescent intensity analyses were performed using Image Studio Lite (Li-Cor). Ephrin WB data were normalized over GAPDH and expressed as median ± interquartile range (IQR). Cell counts, surface area, optical density and volumetric analyses were performed using Fiji (NIH; 1.52i) as previously described [10]. For stripe assays, a 500μm^2^ grid was superimposed over 10 non-overlapping photomicrographs (n=6). GFAP+ astrocytes with nuclei completely inside or outside stripe boundary were counted and expressed as median % GFAP+ nuclei on stripes over total cells counted ± IQR. For proliferation assays, GFAP+/ BrdU+ cells were counted over 10 non-overlapping photomicrographs (n=9) and expressed as ratio over total cell numbers. Data was presented as median BrdU+ astrocyte ratio ± IQR. For wound closure assay, A 5000μm^2^ grid was superimposed over 3 non-overlapping photomicrographs comprising the scratch (n=6). The grid boxes encompassing the scratch was counted at t0 and t48 to obtain initial and final wound surface area, respectively. Data was expressed as median % wound closure at 48h ± IQR. For cFOs and GAP43 cell counts, a 0.05mm^2^ and 0.1mm^2^ grid was superimposed over 5 non-overlapping photomicrographs within 1mm of the infarct core (n=6/ cohort). Double-immunopositive cells were counted and expressed as median double immunopositive cell density ± IQR. For CSPG immunopositive surface area analysis, a 0.1mm^2^ grid was superimposed over 3 adjacent photomicrographs encompassing the entire marmoset V1. A binary mask was generated and thresholded to the intensity of the unaffected rostral calcarine region. Grid boxes encompassing CSPG signal above the threshold were counted. Data was expressed as median CSPG surface area (mm^2^) ± IQR. For optical density analysis, parallel guide lines 200μm apart, starting at the middle of the infarct core and perpendicular to the white matter were superimposed over 3 adjacent GFAP+ or NeuN+ tile scanned photomicrographs. A rectangular marquee was drawn between each guide lines from the white matter to the edge of residual tissue and the average signal intensity was measured within the marquee up to 2000μm and expressed as a percentage over uninjured control. Data was presented as median % GFAP+ optical density ± IQR. Volumetric analysis of the GFAP+ glial scar and NeuN+ residual volume was performed using the Cavalieri method on a full series of GFAP+ or NeuN+ immunolabeled sections (40μm thick, 200μm separation). Sections were imaged under constant exposure settings, as previously described [19]. A binary mask was generated from immunofluorescent photomicrographs and thresholded to visualize the area of dense GFAP+ or NeuN+ labelling. The volume was calculated using a grid (2.5 ×10^5^μm^2^) superimposed on the mask. Results were corrected for linear shrinkage and expressed as median volume (mm^3^) ± IQR.

### Gene expression analysis

Data for the expression of src as well as epha2 and epha4 on human vs. mouse astrocytes and microglia, respectively were extracted from RNAseq data published in [52] (NCBI GEO; GSE73721). Only data from adults (human: >21y;) were used. Data for the expression of efna1 and efna5 on normal, ‘A1’ and ‘A2’ reactive astrocytes in rodents were extracted from microarray gene expression analysis published in [13] (NCBI GEO; GSE35338).

### Statistics and data analysis

The sample size for the *in vivo* study was calculated from preliminary data demonstrating a 20% reduction in glial scar volume following rh-ephrin-A1-Fc treatment. Power analysis for a non-parametric Mann-Whitney U test was conducted using G*Power 3.1 software to determine sufficient sample size using an alpha of 0.05, a power of 0.80, a large effect size (d=2.5), and two tails. Based on these assumptions and equal distribution of samples in each treatment cohorts, the desired sample size calculated was n=6.

No data was excluded during the data analysis for this study. *In vivo* experiments (including all uninjured, injured, vehicle or treatment conditions) were replicated using separate animals. *In vitro* data were replicated (n=6) via separate independent experiments. qPCR, western blots and in-cell-western experiments were replicated using lysates from separate independent experiments with n>3 internal replications per experiment.

In vitro experiments were allocated into treatment conditions using a random number generator. Allocation of in vivo experiments into treatment cohorts was not randomized to enable equal allocation of animal sex between groups. Additionally, in vivo experiments were staggered over 2 years which did not allow for efficient randomization.

The investigator was blinded to treatment cohorts for in vitro data analysis. Since the primary investigator was the main experimenter responsible for animal welfare during the experimental period, data acquisition and analysis the primary investigator was not blinded to the treatment conditions of in vivo experiments.

Statistical calculations were performed using Prism 7 (GraphPad). All measurements presented were from distinct samples and not repeated measures. Based on the D’Agostino or Shapiro Wilk normality test, non-parametric statistics were used in this study, unless otherwise stated. The use of non-parametric statistics in this study offers more robust statistical calculations in experiments with lower sample sizes. Data is not required to fit Gaussian distribution and does not rely on numbers or mean comparison, rather relying on rank-based tests to determine statistical significance between groups, with respect to medians. Histograms are therefore presented as median ± interquartile ranges to reflect the nonparametric statistics used. The non-parametric analysis of variance (ANOVA; Kruskal-Wallis test; KW; 95% CI; unmatched) was performed to determine statistical significance among >2 groups followed by the Dunnet multiple comparisons posthoc test (uncorrected). The non-parametric Mann-Whitney U test (MW; 95% CI; two-tailed; unpaired) was used to determine statistical significance between two groups.

## Acknowledgements

The authors would like to acknowledge de Souza M. and Chan AL. for technical assistance as well as Kania A. and Brotchie A. for manuscript critique. This work was supported by the National Health and Medical Research Council Project Grant (APP20140228) and Senior Research Fellowship to JAB, and the Australian Research Council SRI Stem Cells Australia. The Australian Regenerative Medicine Institute is supported by grants from the State Government of Victoria and the Australian Government.

## Author Contributions

LT and JB designed research. LT, AGB, JHL, ICM and WK performed research. LT and JB analyzed data. LT and JB prepared manuscript.

