## Supplementary Figures for "Replicating infant astrocyte behavior in the adult after brain injury improves outcomes"

### Supplementary Figure 1

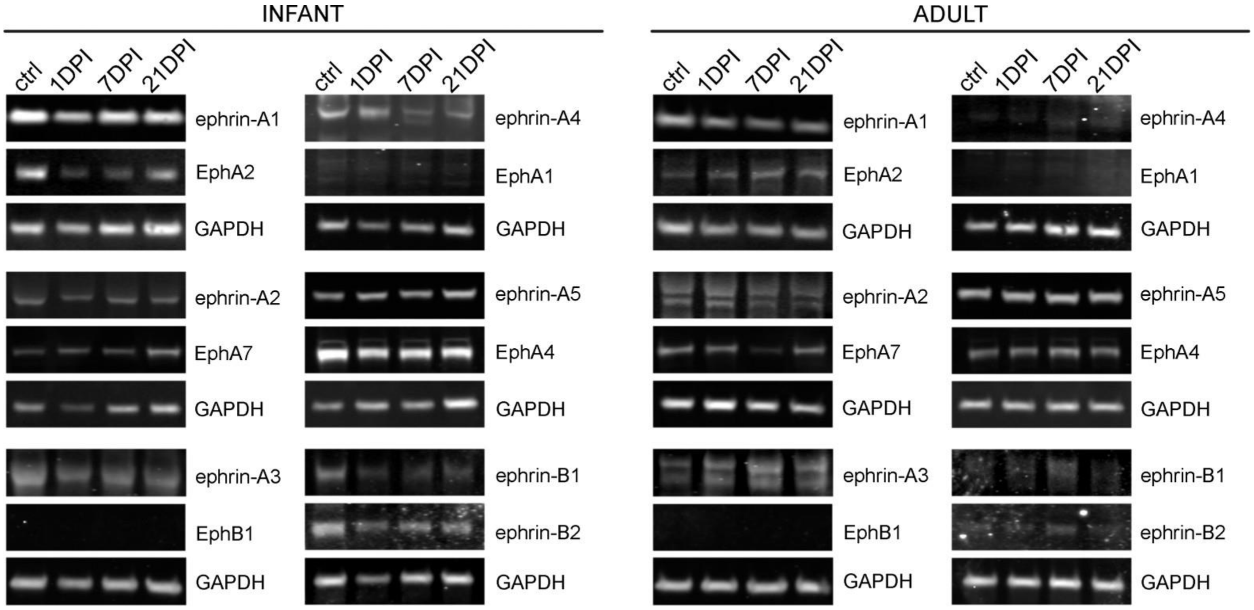


**Supplementary Figure 1. Ephs/ ephrins expression in the infant and adult marmoset V1 post injury.**

Western blot images of ephrin-A1, -A2, -A4, -A4, A5, -B1, -B2 and EphA1, A2, A4, A7, B1 expression in the uninjured infant and adult marmoset V1 as well as at 1, 7 and 21 days post injury (DPI). Result of the densitometry analysis is summarized in Figure 1E.

**Supplementary Figure 2**

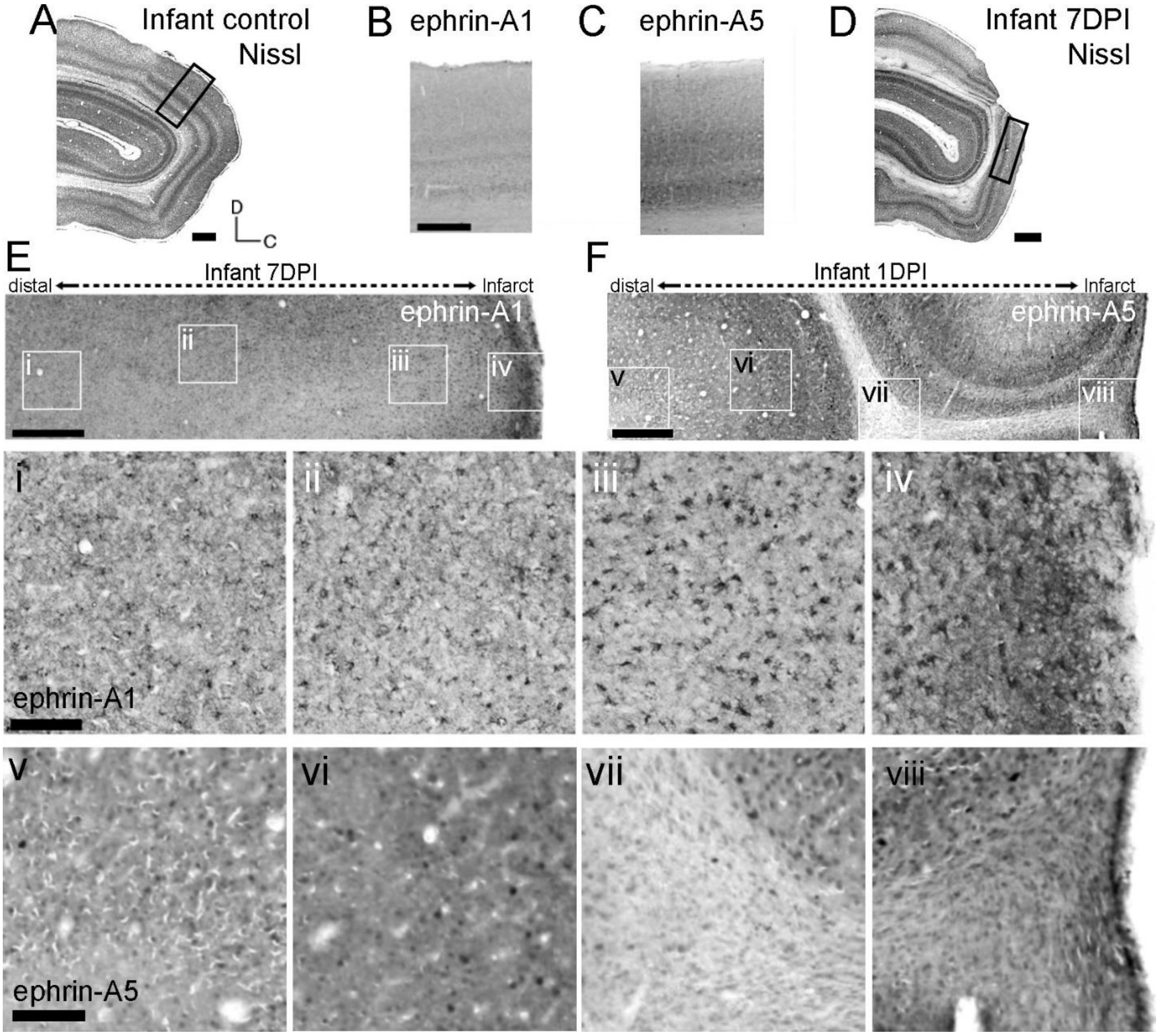


**Supplementary Figure 2. Ephrin-A5 and –A1 is upregulated on Reactive astrocytes in the injured infant V1 at 1 and 7 DPI, respectively.**

(A) Parasagittal Nissl substance-stained section of the control infant V1. Bounding box denotes enlarged regions in (B, C). (B, C) Brightfield photomicrograph ephrin-A1 (B) and –A5 (C) immunolabeled uninjured infant V1. (D) Parasagittal Nissl substance-stained section of the injured infant V1 at 7 DPI. Bounding box denotes enlarged regions in (E, F). (E, F) Brightfield photomicrograph of ephrin-A1 (E) and –A5 (F) immunolabeled section comprising the lesion core and peri-infarct area with bounding boxes enlarged in (i-iv; ephrin-A1) and (v-viii; ephrin-A5) demonstrating dense cellular labelling proximal to the lesion core, diminishing distally. * $p<0.05$. Scale bar: (A, D) 1mm**;** (B, C, E, F) 0.5mm; (i-viii) 200µm.

**Supplementary Figure 3**

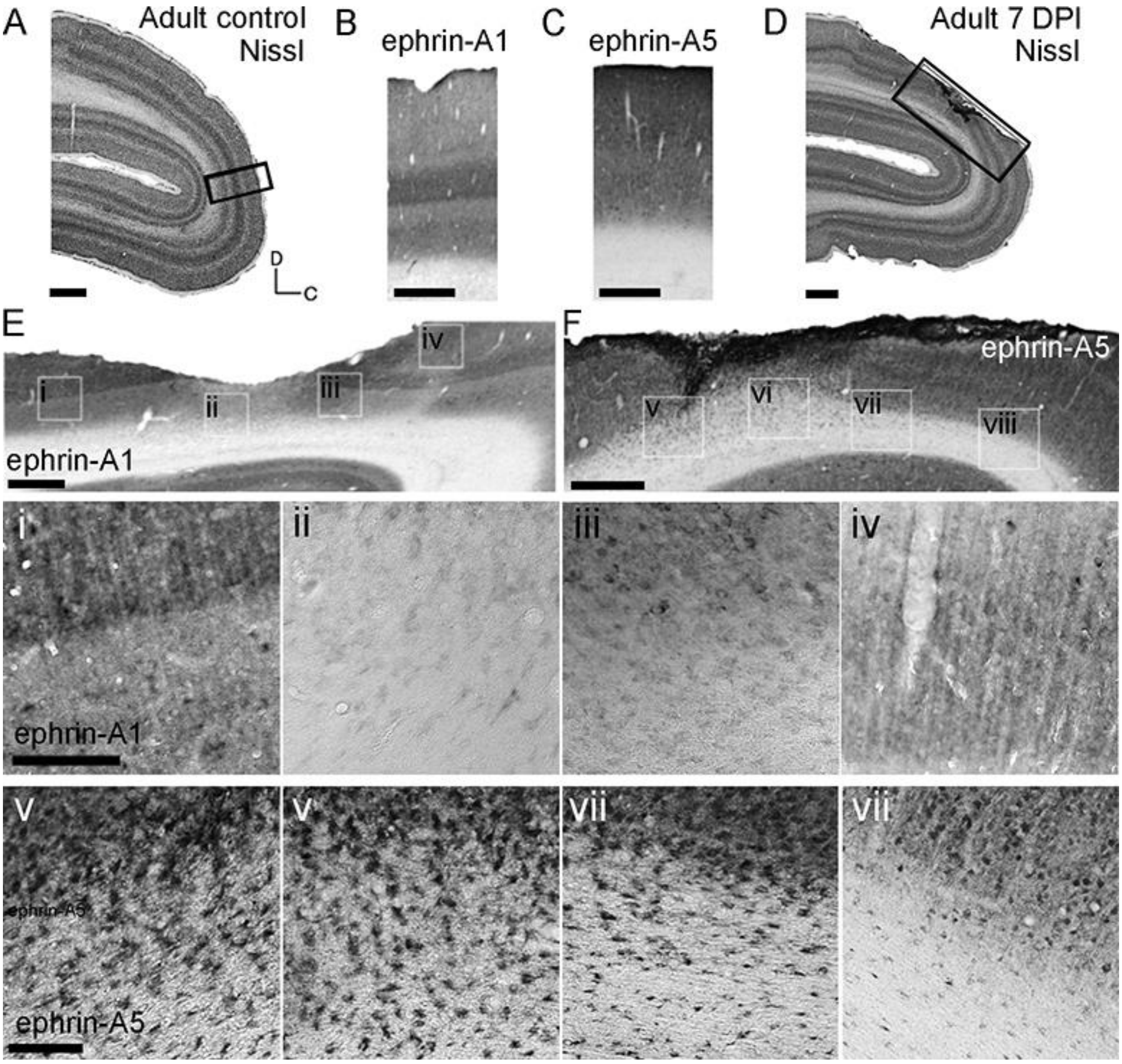


**Supplementary Figure 3. Ephrin-A5 is upregulated on Reactive astrocytes in the injured adult V1 at 1 and 7 DPI.**

(A) Parasagittal Nissl substance-stained section of the control adult V1. Bounding box denotes enlarged regions in (B, C). (B, C) Brightfield photomicrograph ephrin-A1 (B) and –A5 (C) immunolabeled uninjured adult V1. (D) Parasagittal Nissl-stained section of the injured adult V1 at 7 DPI. Bounding box denotes enlarged regions in (E, F). (E, F) Brightfield photomicrograph of ephrin-A1 (E) and –A5 (F) immunolabeled section comprising the lesion core and peri-infarct area with bounding boxes enlarged in (i-iv; ephrin-A1) and (v-viii; ephrin-A5) demonstrating dense cellular labelling of ephrin-A5 proximal to the lesion core, diminishing distally, but not ephrin-A1. * $p<0.05$. Scale bar: (A, D) 1mm**;** (B, C, E, F) 0.5mm; (i-viii) 200µm.

**Supplementary Figure 4**

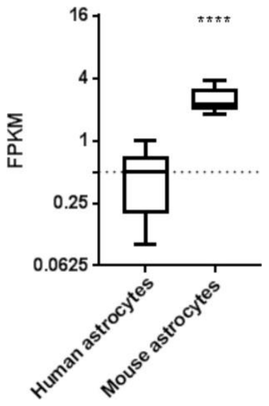


**Supplementary Figure 4. Gene expression analysis demonstrates transcriptional differences that underpins functional differences in reactive astrocyte between rodents and primates**

Expression of Src (c-Src; rodent homolog: pp60c-Src) non-receptor tyrosine kinase was detected at significantly higher levels in adult mouse astrocytes compared to the subthreshold levels detected in adult human astrocytes. This likely contribute to ephrin-A5 induced attraction on human astrocytes. Expression thresholds are as defined in [44]. Database: NCBI GEO GSE73721.

**Supplementary Figure 5**

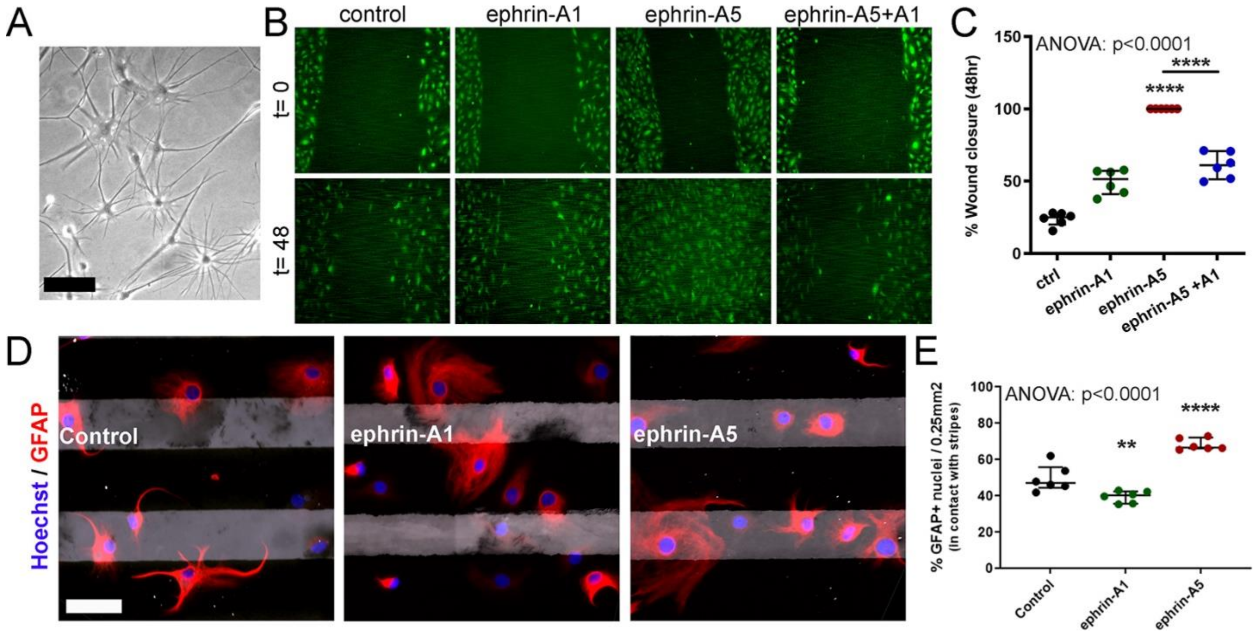


**Supplementary Figure 5. Ephrin-A1 and -A5 treatment induces different outcomes on marmoset astrocyte centripetal migration and guidance.**

(A) Marmoset neurosphere-derived astrocytes in culture under brightfield microscopy. (B, C) in scratch wound assays, ephrin-A5 signalling significantly increased astrocyte wound closure at t=48h while ephrin-A1 signalling did not. Addition of ephrin-A1 significantly attenuated ephrin-A5 induced astrocyte wound closure. (D, E) In stripe assays, ephrin-A5 stripes induced attraction while ephrin-A1 stripes induced repulsion on astrocytes migration. (**) $p<0.01$; (****) $p<0.0001$). Scale: (A) 20µm, (D) 50µm.

**Supplementary Figure 6**

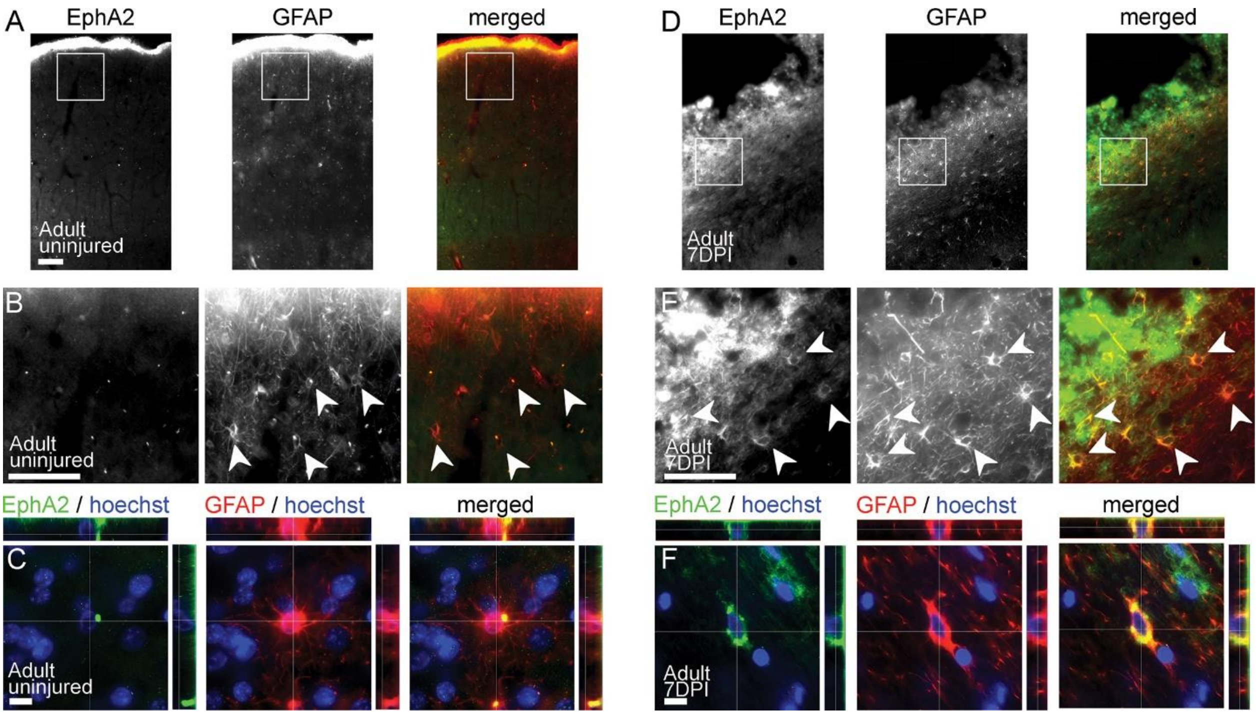


**Supplementary Figure 6. EphA2 is expressed on a subpopulation of reactive astrocytes proximal to the lesion core at 7DPI in adults.**

(A, D) Low magnification EphA2 and GFAP immunofluorescent photomicrographs of uninjured (A) and injured (D) V1. Bounding boxes denotes region enlarged in (B) and (E), respectively. (B, C) EphA2 is not associated with GFAP+ astrocytes in the uninjured marmoset V1. (E, F) EphA2 is associated on a subpopulation of GFAP+ reactive astrocytes proximal to the injury site in the injured adult V1, at 7DPI. Scale bar: (A, D): 1mm; (B, E): 0.5mm; (C, F): 20µm.

**Supplementary Figure 7**

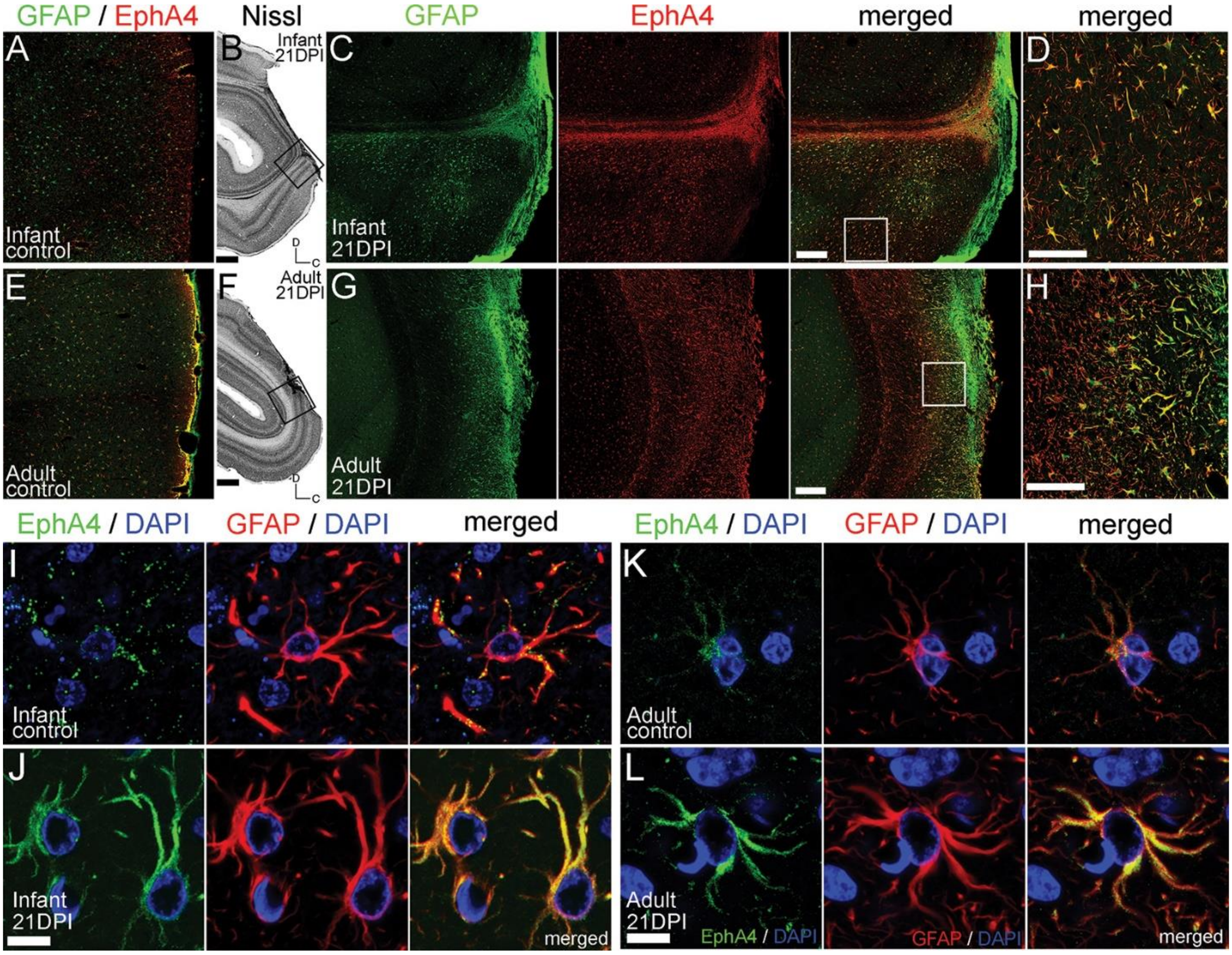


**Supplementary Figure 7. EphA4 is expressed on a subpopulation of reactive astrocytes proximal to the lesion core in the injured infant and adult marmoset V1.**

(A, E) Low magnification EphA4 and GFAP immunofluorescent photomicrographs of uninjured) infant (A) and adult (E) V1. (B, C) Nissl substance histology stained parasagittal sections of the injured infant (B) and adult (F) V1 at 21 DPI with bounding boxes denoting regions enlarged in (C) and (G), respectively. (C, G) Low magnification EphA4 and GFAP immunofluorescent photomicrographs of the injured infant (C) and adult (G) V1. Bounding boxes denotes region enlarged in (D) and (H), respectively. Minimal EphA4 expression was detected on GFAP+ astrocytes in the uninjured infant (I) or adult (K) V1. (C, D, G, H) EphA4 expression was increased on a subpopulation on GFAP+ reactive astrocytes proximal to the injury site in infants (C, D, J) and adults (G, H, L) at the peak of EphA4 upregulation (Fig. 1E). Scale bar: (A, C, E, G): 0.5mm; (B, F): 1mm; (D, H): 100µm; (I-L): 20µm.

**Supplementary Figure 8**

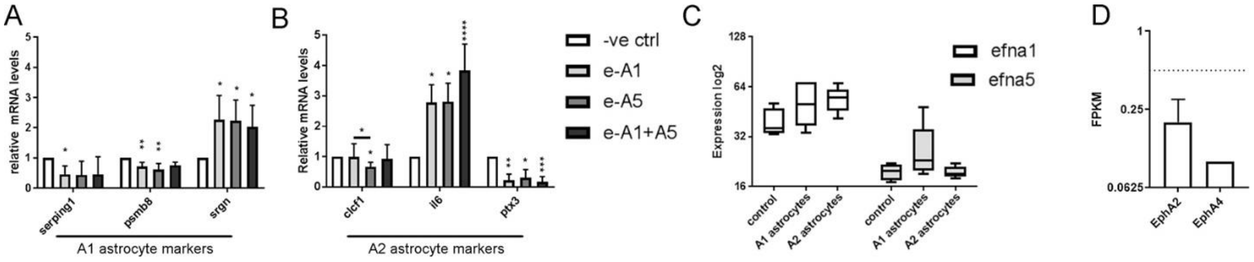


**Supplementary Figure 8. Ephrin-A1 or -A5 signalling does not drive reactive astrocyte phenotype changes into either A1 or A2 subtypes.**

(A, B) qRT-PCR analysis of A1 (A) and A2 (B) reactive astrocyte subtype markers (as defined in **), following ephrin-A1 and/or -A5 signalling on IL6 stimulated human Reactive astrocytes. Neither ephrin-A1 nor ephrin-A5 individually or combined was responsible for- or sufficient to induce the conversion of IL6 stimulated human Reactive astrocytes into either A1 or A2 reactive astrocyte subtypes. (C) Transcriptional profiling revealed that neither A1 nor A2 reactive astrocyte subtypes derived from mice upregulates ephrin-A1 or –A5. (D) Subthreshold levels of EphA2 and EphA4 were detected on human microglia. Absence of either Eph receptor indicate the negligible local immune response from rh-ephrin-A1-Fc infusion. * $p<0.05$; ** $p<0.01$; *** $p<0.001$; ****$p<0.0001$. (A-C) Database: NCBI GEO GSE35338 [5]. (D) Expression thresholds are as defined in [44]. Database: NCBI GEO GSE73721.

**Supplementary Figure 9**

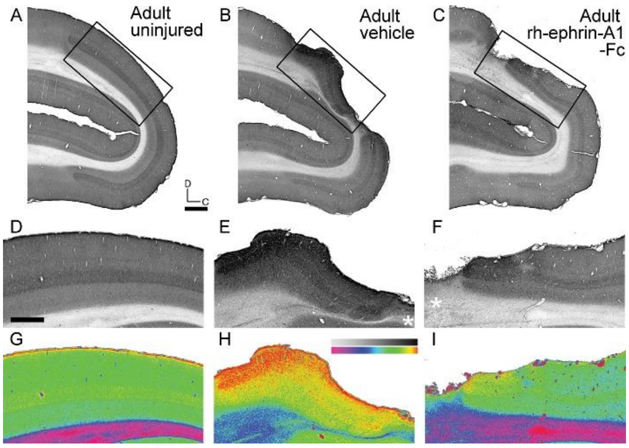


**Supplementary Figure 9. Rh-ephrin-A1-Fc treatment markedly reduces chondroitin sulfate proteoglycan (CSPG) deposition in the gliali scar after injury**

(A-C) Low magnification parasagittal sections of the adult marmoset V1 immunolabeled for chondroitin sulfate proteoglycans (CSPGs). Bounding boxes denote enlarged regions in (D-F), respectively. (*) indicates lesion core. To aid visual assessment of CSPG labelling intensity, images were normalised against the distal, ventral white matter, and a pre-set, artificial look-up-table (ImageJ, Spectrum) was applied to (D-F) to generate pseudo-coloured heatmap images in (G-I), respectively. Rh-ephrin-A1-Fc treatment (I) resulted in a marked decrease in CSPG deposition proximal to the lesion core, compared to vehicle control (H). Scale bar: (A-C): 1mm; (D-F): 0.5mm.

**Supplementary Figure 10**

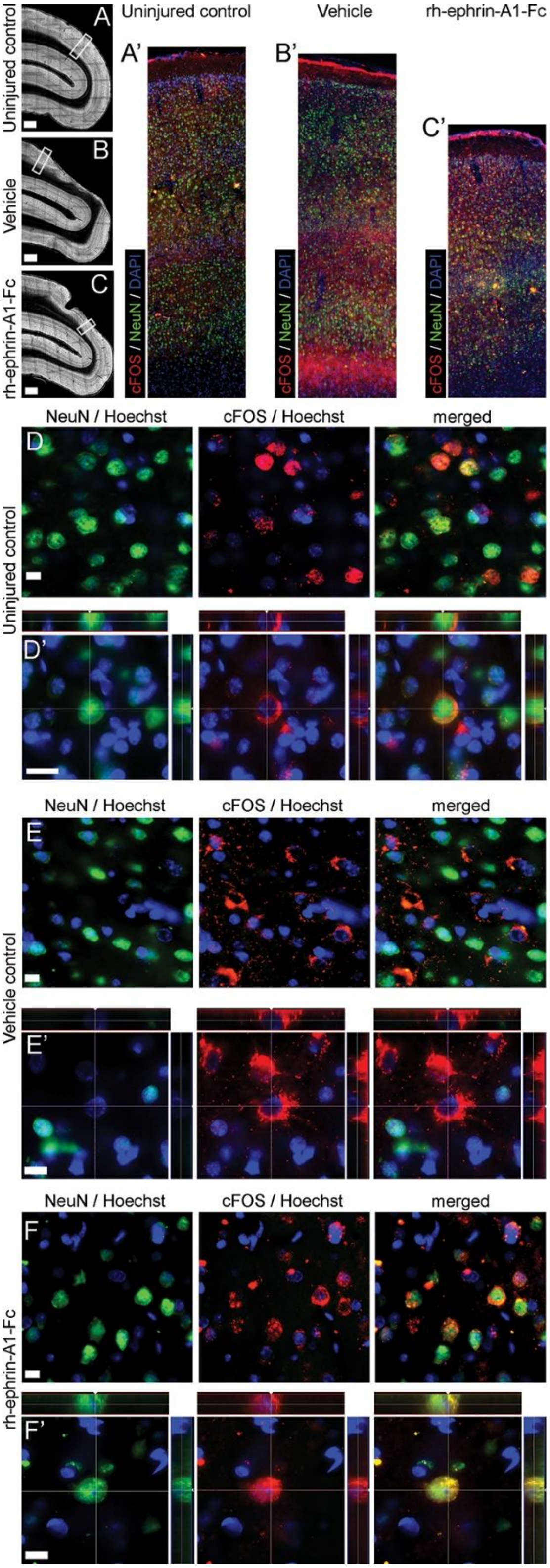


**Supplementary Figure 10. cFOS expression on neurons is reduced following injury but is rescued by rh-ephrin-A1-Fc infusion.**

(A-C) DAPI stained fluorescent parasagittal section of the uninjured (A), vehicle treated (B) and rh-ephrin-A1-Fc treated (C) adult marmoset V1. Bounding boxes correspond to enlarged NeuN and cFOS immunofluorescent labelled photomicrographs within 1mm of infarct epicenter in (A'-C'). (D, D') In uninjured controls, and rh-ephrin-A1-Fc treated conditions (F, F'), cFOS expression (red) is primarily associated with NeuN+ neurons (green). (E, E') In vehicle controls a substantial subpopulation of cFOS+/ NeuN- cells was observed primarily in the white matter (B'). Scale bar: (A-C) 1mm; (D-F) 20μm; (D'-F') 10μm.

**Supplementary Figure 11**

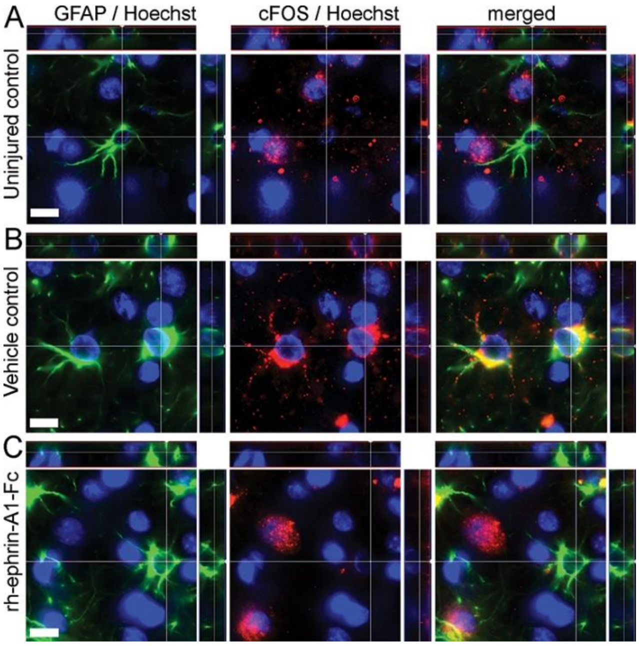


**Supplementary Figure 11. cFOS expression is associated with a subpopulation of GFAP+ reactive astrocyte proximal to the lesion core but is absent after rh-ephrin-A1-Fc infusion.**

(A) cFOS expression (red) is not associated with GFAP+ (green) reactive astrocyte in uninjured (A) or rh-ephrin-A1-Fc treated (C) conditions. (D) The subset of cFOS+/NeuN- cells identified in Supplementary Figure 6E, E' is identified to be a subpopulation of cFOS+/GFAP+ reactive astrocyte present proximal to the lesion core. Expression of cFOS on Reactive astrocytes is associated with increased proliferation. Scale bar: (A-C) 10µm.

**Supplementary Figure 12**

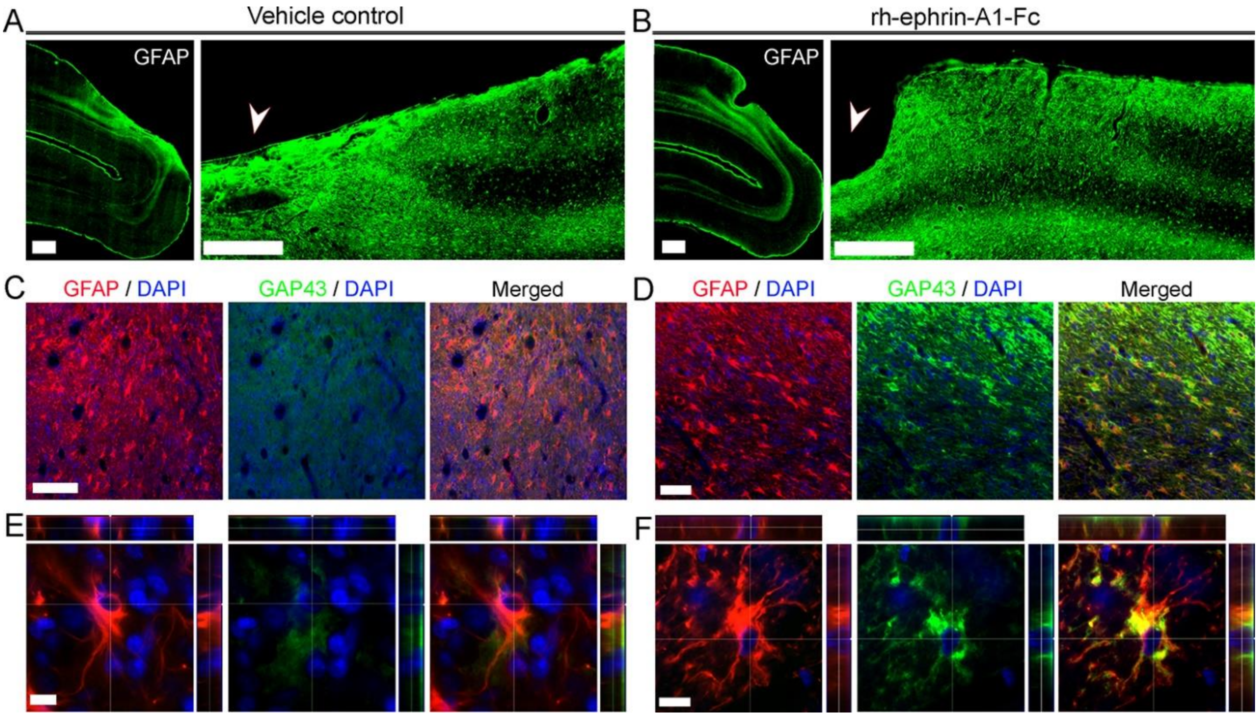


**Supplementary Figure 12. Rh-ephrin-A1-Fc infusion induced increased GAP43 expression on GFAP+ reactive astrocytes proximal to the lesion site.**

(A, B) GFAP immunofluorescent labelled parasagittal sections of vehicle (A) and rh-ephrin-A1-Fc (B) treated V1 with enlarged lesion core (arrow) and peri-infarct area. (C, D) High magnification photomicrographs of GFAP and GAP43 immunofluorescent labelled sections within 1mm proximal to the injury core (arrow: A, B). (C, E) GAP43 expression is not associated with GFAP+ Reactive astrocytes in vehicle treated controls. (D, F) Following rh-ephrin-A1-Fc infusion, a considerable population of GAP43+/GFAP+ Reactive astrocytes were detected proximal to the injury core. Scale bar: (A, B) 1mm; (C, D) 100µm; (E, F) 10µm.
